# Soil, senescence and exudate utilisation: Characterisation of the Paragon var. spring bread wheat root microbiome

**DOI:** 10.1101/2021.02.09.430461

**Authors:** Sam Prudence, Jake Newitt, Sarah F. Worsley, Michael C. Macey, J. Colin Murrell, Laura E. Lehtovirta-Morley, Matthew I. Hutchings

**Affiliations:** Department of Molecular Microbiology, John Innes Centre, Norwich Research Park, Norwich. NR47UH; School of Biological Sciences, University of East Anglia, Norwich Research Park, Norwich. NR4 7TJ; School of Environmental Sciences, University of East Anglia, Norwich Research Park, Norwich. NR4 7TJ

## Abstract

Conventional methods of agricultural pest control and crop fertilisation are contributing to a crisis of biodiversity loss, biogeochemical cycle dysregulation, and ecosystem collapse. Thus, we must find ecologically responsible means to control disease and promote crop yields. The root-associated microbiome may contribute to this goal as microbes can aid plants with disease suppression, abiotic stress relief, and nutrient bioavailability. We applied 16S rRNA gene & fungal 18S rRNA gene (ITS2 region) amplicon sequencing to profile the diversity of the bacterial, archaeal & fungal communities associated with the roots of UK elite spring bread wheat variety *Triticum aestivum var.* Paragon in different soils and developmental stages. This revealed that community composition shifted significantly for all three groups across compartments. This shift was most pronounced for bacteria and fungi, while we observed weaker selection on the ammonia oxidising archaea-dominated archaeal community. Across multiple soil types we found that soil inoculum was a significant driver of endosphere community composition, however several bacterial families were identified as core enriched taxa in all soil conditions. The most abundant of these were *Streptomycetaceae* and *Burkholderiaceae.* Moreover, as the plants senesce, both families were reduced in abundance, indicating that input from the living plant was required to maintain their abundance in the endosphere. To understand which microbes are using wheat root exudates in the rhizosphere, root exudates were labelled in a ^13^CO_2_ DNA stable isotope probing experiment. This shows that bacterial taxa within the *Burkholderiaceae* family among other core enriched taxa, such as *Pseudomonadaceae,* were able to use root exudates but *Streptomycetaceae* were not. Overall, this work provides a better understanding of the wheat microbiome, including the endosphere community. Understanding crop microbiome formation will contribute to ecologically responsible methods for yield improvement and biocontrol in the future.

## Introduction

Wheat is a staple crop for more than 4 billion people and globally accounts for more than 20% of human calorie and protein consumption [1]. This means that farming wheat, and the accompanying use of chemical fertilisers and pesticides, has a huge environmental impact worldwide. For example, up to 70% of nitrogen fertiliser is lost each year through runoff and microbial denitrification which generates the potent greenhouse gas N_2_O [2]. The challenge facing humans this century is to grow enough wheat to feed an increasing global human population while reducing our reliance on agrochemicals which contribute to climate change and damage ecosystems [3]. One possible way to achieve this is to manipulate the microbial communities associated with wheat and other crop plants. These communities are commonly referred to as “microbiomes” and a healthy microbiome can enhance host fitness by providing essential nutrients [4], increasing resilience to abiotic stressors [5], and protecting against disease [4]. Each new generation of plants must recruit the microbial species (archaea, bacteria, fungi and other micro-eukarya) that make up its root microbiome from the surrounding soil, and this means that the soil microbial community is an important determinant of plant root microbiome composition [6].

Plants are able to influence the microbial community in the rhizosphere, which is the soil most closely associated with the roots, and the endosphere, which is the inside of the roots. The microbes within these root-associated environments tend to have traits which benefit the host plant [7] and plants modulate these microbial communities by depositing photosynthetically fixed carbon into the rhizosphere in the form of root exudates, a complex mixture of organic compounds consisting primarily of sugars, organic acids, and fatty acids [8]. Plants deposit up to 40% of their fixed carbon into the soil [9], and there is evidence to suggest that certain molecules within these exudates can attract specific bacterial taxa [6,10,11]. Thus, the implication is that host plants attract specific microbial taxa from a diverse microbial soil community, and generate a root microbiome that contains only the subset of the soil community most likely to offer benefits to the host plant [12]. In return, the growth of beneficial microbes is supported by the nutrients from root exudates, such that the plants and microbes exchange resources in a mutually beneficial symbiosis. Traditional plant breeding may have had a negative effect on this process in important food crops such as barley and wheat; for example, selection for traits such as increased growth and yield may have inadvertently had a negative influence on root exudation and microbiome formation [8,11]. Long-term use of fertilizer also reduces the dependency of the host plant on microbial interactions, further weakening the selective pressure to maintain costly exudation of root metabolites [13]. This highlights the need for a greater understanding of the factors that underpin microbiome assembly and function in important domesticated crop species such as bread wheat, *Triticum aestivum*.

To understand the key functions in a host-associated microbial community, it can be useful to define the core microbiome, i.e. the microbial taxa consistently associated with a particular plant species regardless of habitat or conditions and which provide a service to the host plant and/or the broader ecosystem [14,15]. The core microbiome of the model plant *Arabidopsis thaliana* is well studied [6,16], and it has also been characterized for numerous other plant species to varying degrees [17–19]. In elucidating the core microbiome a number of factors must be accounted for, including soil type [20,21], developmental stage [22,23], genotype [8,22,24] and, in the case of crop plants, agricultural management strategy [23,25–27]. The core microbiome has been investigated for bread wheat [28–32] and, while most studies focus on the rhizosphere, Kuźniar *et al.* [28] identified a number of core bacterial genera within the endosphere including *Pseudomonas* and *Flavobacterium*. Their study focussed on a single soil type and developmental stage, but to reliably identify the core microbial taxa associated with wheat, more of the aforementioned factors must be analysed. Microbial community surveys are also often limited to investigations of bacterial or, in some cases, fungal diversity meaning that knowledge of wheat root community diversity is limited to these two groups. Root-associated archaea are considerably understudied, particularly within terrestrial plant species such as wheat. Most generic and commonly used 16S rRNA gene PCR primer sets fail to capture archaeal diversity [33], thus the diversity of archaea within soils is commonly overlooked. Key soil groups such as ammonia oxidizing archaea (AOA) play a significant role in nitrogen cycling, a key ecological service, and one study has managed to link an AOA to plant beneficial traits [34], suggesting that the role of archaea within the terrestrial root associated microbiome warrants further study.

For many important crops such as wheat, barley, maize, corn, and rice, developmental senescence is a crucial determinant of yield and nutrient content [35,36]. Developmental senescence occurs at the end of the life cycle, and during this process, resources, particularly nitrogen, are diverted from plant tissues into the developing grain [35,36]. Senescence represents a dramatic shift in the metabolic activity of the plant [35] and in the regulation of pathways of pathogen defence [36,37]. Given that root exudation is a dynamic process [38], it would be reasonable to assume that senescence affects root exudation substantially, particularly because of the diversion of nitrogen to the developing grain (several major wheat root exudate compounds, like amino acids, nucleosides, and numerous organic acids, contain nitrogen [38]). To our knowledge, changes within the wheat root microbial community during wheat senescence have not been investigated previously. Given the pivotal role senescence plays in grain development and yields, microbial community dynamics during this process warrant investigation. At the onset of senescence, plant resources are redirected to the seed, root exudation is reduced, and root tissues start to decay. It is plausible that this shift in plant metabolism would cause a change in the root-associated microbiome, and greater understanding of this could come inform agricultural management strategies and the design of new crop cultivars.

One major limitation of metabarcoding approaches is that they do not reveal which microbial taxa are actively interacting with plants, for example via the utilisation of compounds exuded by the roots. ^13^CO_2_ DNA stable Isotope Probing (SIP) is a powerful tool for investigating the role of root exudates in microbiome assembly. As plants are incubated with ^13^CO_2_, the heavy carbon is fixed and incorporated into exuded organic compounds. Microbial communities that actively metabolise root exudates will incorporate ^13^C into their DNA and can thus be identified [9,39]. While numerous DNA-SIP studies have probed metabolically active communities associated with wheat, few have assessed root exudate metabolism directly using high-throughput sequencing methods for microbial identification [40,41]. Of the two studies that have, similar findings were presented but with some distinct differences. Both studies showed that exudate-metabolising microbial communities in the rhizosphere consisted primarily of Actinobacteria and Proteobacteria [42,43]. Taxa from Burkholderiales specifically were shown to dominate exudate metabolism in one study [42], whereas the other highlighted *Paenibacillaceae* as exudate metabolisers within the rhizosphere [43]. Discrepancies between these studies likely result from different soil types and wheat genotypes, and this demonstrates a need for further DNA-SIP experiments using different soils and different wheat varieties.

In this study we characterised the rhizosphere and endosphere microbiomes of *Triticum aestivum* variety Paragon, an UK elite spring bread wheat, using metabarcoding and ^13^CO_2_ DNA-SIP. Although wheat rhizosphere bacterial communities have been well characterised under a wide range of conditions [22,24,29–31,44], few studies have surveyed the endosphere community. Here, we profile the archaeal, bacterial and fungal communities in the bulk soil, rhizosphere and endosphere compartments of *T. aestivum* var. Paragon using 16S rRNA gene and ITS2 amplicon sequencing. We further characterise the bacterial communities using ^13^CO_2_ DNA-SIP. We aimed to address the following questions: (1) Are there any core microbial taxa within the endosphere and rhizosphere of *T. aestivum var.* Paragon across starkly contrasting soil environments? (2) How does the community change as the plant enters developmental senescence, and which microbial taxa, if any, are unable to persist through senescence? (3) Do wheat roots select for specific archaeal lineages as they do for bacteria and fungi? (4) Which bacterial taxa utilise wheat root exudates? The results provide a significant advance towards understanding wheat-microbiome interactions and establishing an understanding of the core microbial taxa in *T. aestivum* var. Paragon.

## Results

### The microbial community associated with *Triticum aestivum* var. Paragon

To gain initial insights into the microbial communities associated with wheat roots, we characterised the microbial community associated with field-grown wheat sampled during the stem elongation growth phase. The diversity of microbes in the bulk soil, rhizosphere, and endosphere compartments was investigated using 16S rRNA gene (for bacteria and archaea) and ITS2 (for fungi) metabarcoding, respectively. The bacterial and fungal communities differed significantly across compartments (bacterial PERMANOVA: R^2^=0.8, *p* < 0.01; fungal PERMANOVA: R^2^=0.63, *p* < 0.01). This was particularly the case for the rhizosphere and endosphere compartments compared to bulk soil, as demonstrated by principal coordinates analysis (PCoA) (Figure 1; A1, A3). Community profiles did not indicate a strong shift in the archaeal community across compartments at the family level (Figure 2; C1), but statistical analysis indicated a significant effect of compartment on archaeal community composition at the OTU level (archaeal PERMANOVA: R^2^=0.66, *p* < 0.01), with PCoA indicating that differences in the endosphere may mostly be responsible for this shift (Figure 1; A2). For the bacterial community, the family *Streptomycetaceae* showed the greatest average relative abundance in the endosphere (25.12%), followed by *Burkholderiaceae* (11.99%) and *Sphingobacteriaceae* (7.75%). In the rhizosphere the relative abundance of *Streptomycetaceae* was much lower (2.58%), while *Micrococcaceae* were most abundant (8.43%), followed by *Burkholderiaceae* (7.41%) and *Sphingobacteriaceae* (6.58%) (Figure 2; A1). The fungal endosphere community was dominated by the Xyariales order (32.9%), followed by the class Sordariomycetes (14.33%), then the *Metarhizium* (10.44%). For the rhizosphere, however *Metarhizium* had the greatest relative abundance (27.36%), followed by the Chaetothyriales order (12.32%) and the Sordariomycetes (9.23%). The archaeal community was overwhelmingly dominated by the AOA family *Nitrososphaeraceae* (endosphere 89.77%, rhizosphere 81.55%). Differential abundance analysis demonstrated that the abundance of fourteen bacterial families, including *Streptomycetaceae*, *Burkholderiaceae* and *Sphingobacteriaceae,* increased significantly within the rhizosphere and/or the endosphere relative to the bulk soil (Figure 2; A, Figure 3; A1). The families *Streptomycetaceae* (16.4% contribution, *p* < 0.01) and *Burkholderiaceae* (6.1% contribution, *p* < 0.01) were the two most significant contributors to the bacterial community shift as confirmed by SIMPER analysis (Supplementary Table 1). For the fungal community, most significantly differentially abundant groups were reduced in abundance compared to in the bulk soil, however one taxon was significantly more abundant in the rhizosphere *(Mortierellaceae),* and one was significantly more abundant in the endosphere *(Parmeliaceae)* (Figure 3; A2). No significantly differentially abundant archaeal families were found.

**Figure 1.**
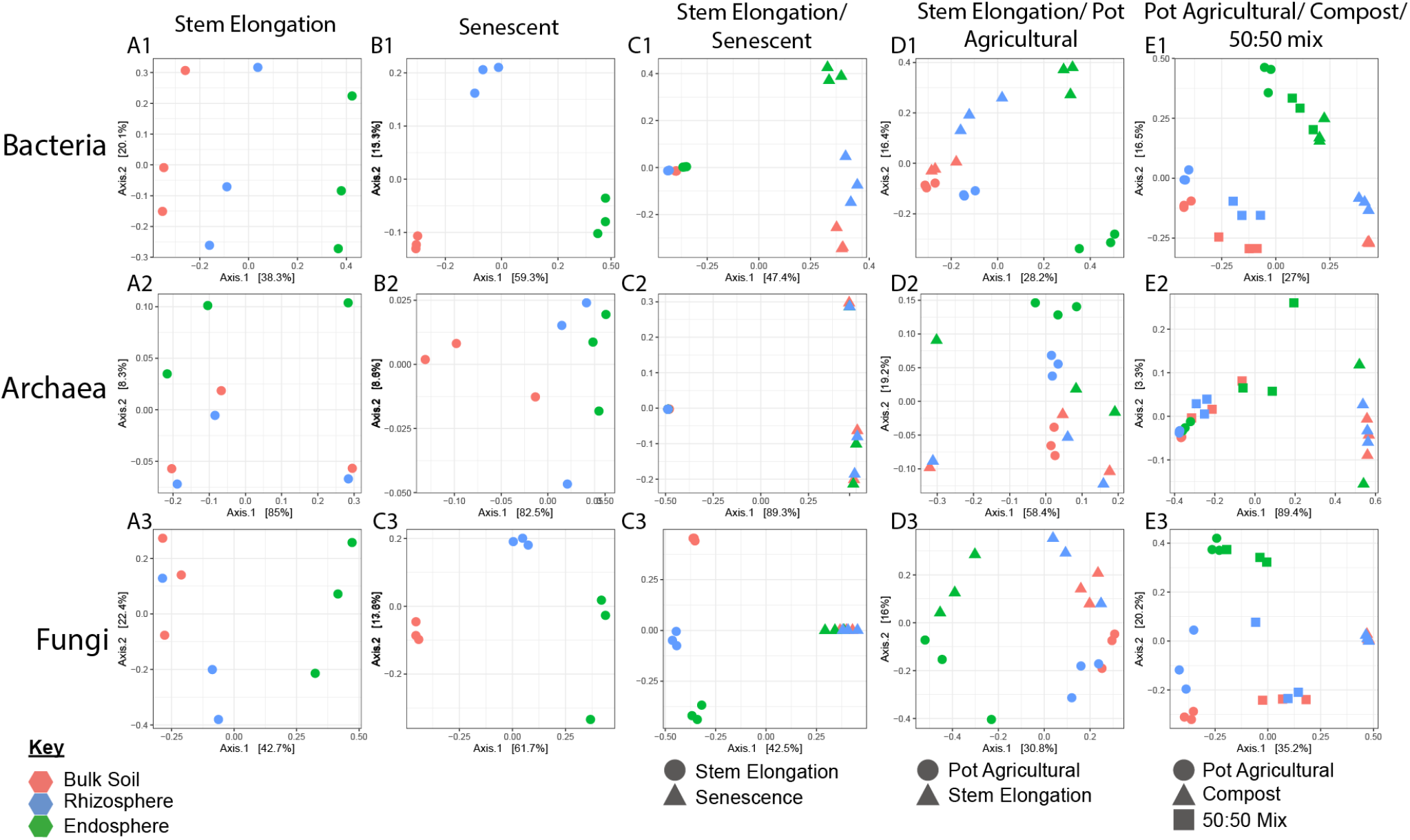
Principal Coordinates Analysis (PCoA) performed on Bray Curtis dissimilarities between samples of the bacterial, archaeal and fungal communities associated with wheat roots. Colours indicate root compartment; green = endosphere, blue = rhizosphere and pink = bulk soil. N=3 replicate plants per treatment. A1, A2, A3 show PCoA for Plants cultivated at the Church Farm field studies site at the stem elongation growth phase. B1, B2 and B3 show data from plants after senescence. C1, C2 and C3 show comparisons between stem elongation growth phase (circles) and senescent plants (triangles). D1, D2 and D3 show comparisons between 4-week-old laboratory cultivated plants (circles) and stem elongation growth phase field cultivated plants (triangles). E1, E2 and E3 show PCoA comparing communities associated with plants cultivated under laboratory conditions in agricultural soil (circles), Levington F2 compost (triangles) or a 50:50 mix of the two (squares).

**Figure 2.**
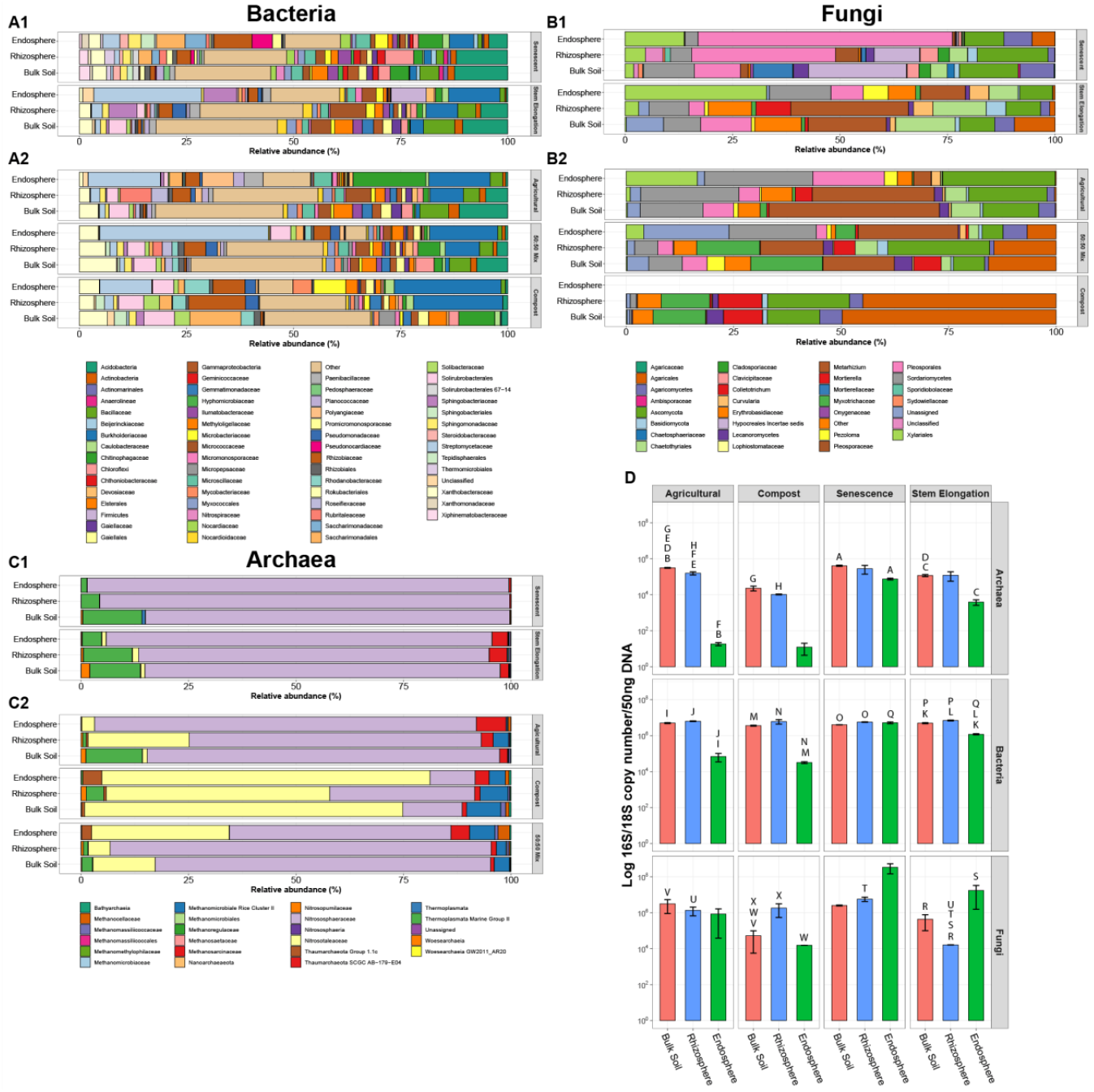
The mean relative abundance (%) of each bacterial, fungal or archaeal taxon within the endosphere, rhizosphere or bulk soil of stem elongation growth phase and senesced wheat plants. Plants were grown at the Church Farm field studies site (**A1, B1, C1**) or under laboratory conditions in agricultural soil, Levington F2 compost or a 50:50 mix of the two (**A2, B2, C2**) (N=3 replicate plants per treatment). Colours indicate different microbial taxa (bacterial, fungal or archaeal). For the archaeal community, N=2 replicate plants for the endosphere of plants grown in Levington F2 compost. Within stacked bars taxa are shown in reverse alphabetical order (left to right). **D**, qPCR data demonstrating the abundance of fungi, bacteria or archaea within the root microbiome. Bars show the mean log 16S or 18S rRNA gene copy per 50 ng of DNA within the endosphere, rhizosphere or bulk soil compartment of plants. Plants were grown in agricultural soil or compost (first and second column, respectively), or were those sampled from the Church farm field studies site during developmental senescence or during the stem elongation growth phase (third and fourth column, respectively). N=3 replicate plants per treatment. Bars represent ± standard error of the mean. Letters indicate a statistically significant difference between the two samples (Tukeys HSD, *p* < 0.05 for all).

**Figure 3.**
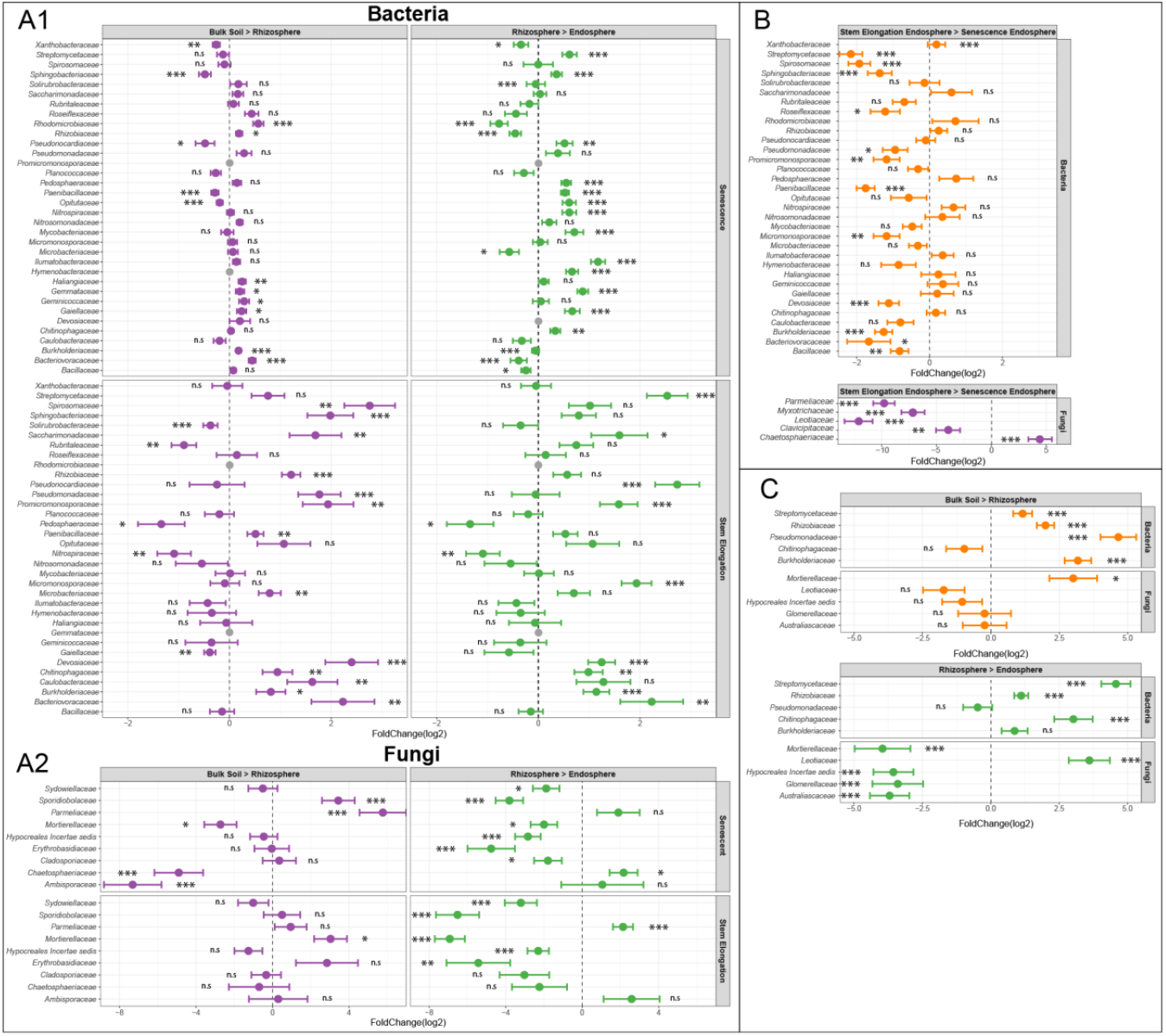
Results of differential abundance analysis. Dots show the log2 fold change of different bacterial or fungal families and error bars show ± log fold change standard error. Results are from N=3 replicate plants per treatment. Shown are: **A** Bacterial and fungal families that were differentially abundant between the bulk soil the rhizosphere, and between the rhizosphere and the endosphere for stem elongation and senesced plants. **B** Bacterial and fungal taxa that were differentially abundant between the endosphere of stem elongation growth phase plants and senesced plants. **C** Bacterial and fungal taxa that were differentially abundant regardless of soil type for pot grown wheat. Analysis was performed using DESeq2. If a family had a base mean > 200 and a significant p-value (significance cut-off *p* < 0.05) in one or more comparison, data for that taxon was plotted for all comparisons, * indicates *p* < 0.05, ** indicates *p* < 0.01, *** indicates *p* < 0.001, and n.s indicates *p* > 0.05. Data for all pot-grown plants were pooled and taxa which still showed significant fold change across compartments were included. For all complete statistical outputs see Supplementary Tables 10-16.

Quantitative PCR (qPCR) was used to estimate the total abundance of archaeal and bacterial 16S rRNA genes and fungal 18S rRNA genes (Figure 2; D). This showed that bacterial 16S rRNA gene copy number was significantly greater within the bulk soil and the rhizosphere compartments when compared to the endosphere (Tukey’s HSD, *p* < 0.01 for both comparisons). Fungi outnumbered bacteria and archaea by more than an order of magnitude within the endosphere (Figure 2; D). This may indicate that fungi are more abundant within the endosphere but could also be a product of the higher 18S rRNA gene copy number per genome within some fungi [45]. When comparing bulk soil to the endosphere, archaeal 16S rRNA gene copy number decreased by two orders of magnitude in the endosphere, while the fungal 18S rRNA gene copy number increased by two orders of magnitude. Despite this, root compartments were not found to significantly influence the abundance of archaea or fungi (ANOVA, *p >* 0.05). This is likely due to high variation across the replicates and could indicate more stochastic root colonisation by fungi and archaea. Compared to bacteria or fungi there were at least three orders of magnitude fewer archaeal 16S rRNA gene copies detected within the endosphere. Despite the lower 16S rRNA gene copy number found in most archaeal genomes [46] this likely demonstrates archaea colonise the root in much lower numbers than the other root microbiota.

### The effect of developmental senescence on the root community

We next aimed to investigate the effect of developmental senescence on the root microbial community and, specifically, to identify microbial taxa associated with the roots of living plants that decline in number during senescence. Developmental senescence is the final stage in wheat development and the point at which nutrients become remobilised from the plant into the developing grain. At this point the plants are no longer green or actively growing. Senescent plants were sampled from the same site as the plants sampled during stem elongation growth phase. Analysis of rRNA gene copy number (from qPCR experiments) showed that plant growth phase significantly influenced the abundance of bacteria (growth phase in a linear model: F-value = 4.86, *p* < 0.05) and archaea (F-value = 10.55, *p* < 0.01 in a linear model) within the root microbiome (Figure 2; D). Comparing specific compartments for each group showed that, while there was no significant difference in the abundance of bacteria within the bulk soil or rhizosphere sampled at either growth phase (Tukey’s HSD, *p* > 0.05), the abundance of bacteria increased significantly within the endosphere after senescence (Tukey’s HSD, *p* < 0.001). Fungal 18S was significantly reduced in the rhizosphere after senescence (Tukey’s HSD, *p* < 0.05) but increased by an order of magnitude in the endosphere, although this increase was not statistically significant (Tukey’s HSD, *p* > 0.05), likely due to variation across replicates. For archaea there were no statistically significant differences in 16S rRNA gene copy number between the two growth phases for any compartment. Both fungal and bacterial community composition differed significantly across the three different root compartments of senescent plants, as clearly demonstrated by PCoA (Figure 1; B1, B2, B3) and PERMANOVA analysis for all three microbial groups (Supplementary Table 2). In addition to this, PCoA showed a clear difference between the microbial communities associated with senescent or stem elongation growth phase plants, however, they also indicated that the root community was much more variable for senescent plants compared to those in the stem elongation phase (Figure 1; C1, C2, C3). PERMANOVA analysis corroborates this observation as, whilst this showed a significant effect of plant growth phase on overall community composition for all three microbial groups (PERMANOVA, bacterial: R^2^=0.47, *p* < 0.001, archaeal: R^2^=0.89, *p* < 0.001, fungal: R^2^=0.42, *p* < 0.001), betadisper analysis indicated that microbial community dispersion was not equal between the two growth phases (*p* < 0.01 for all), i.e. the senescent growth phase showed greater community variability compared to the stem elongation phase.

For individual taxa, differential abundance analysis showed that sixteen bacterial and fungal taxa were significantly less abundant within the endosphere of senesced plants than at the stem elongation growth phase (*p* < 0.05, Supplementary Table 14). The largest change in abundance was a two-fold reduction in the family *Streptomycetaceae* and there was also a significant reduction in the relative abundance of the families *Burkholderiaceae* and *Sphingobacteriaceae* in senescent plants (Figure 3; A1, B). This implies that these taxa may require input from the living plant in order to persist within the endosphere. No archaeal taxa demonstrated significant changes in abundance across root compartments between growth phases. The archaeal community was consistently dominated by the AOA family *Nitrososphaeraceae*. For the fungal community, differential abundance analysis indicated that the abundance of most taxa was significantly reduced in senescent plants, with the exception of *Chaetosphaeriaceae* which showed a four-fold increase during senescence when compared to the stem elongation phase.

### Laboratory-grown *Triticum aestivum* var. Paragon plants provide an agriculturally relevant model

Root associated microbial communities can be influenced by a multitude of abiotic factors, including crop cultivation practices and climatic conditions [47]. To test whether the microbiomes of laboratory-grown plants are comparable to those grown in the field, plants were grown for four weeks under laboratory conditions in soil collected from the Church Farm site and the composition of the root microbiome was profiled using 16S rRNA gene and ITS2 metabarcoding. Laboratory-grown plants were sampled during root growth phase, whereas field plants were sampled during the late stem elongation growth phase, meaning laboratory-grown plants were sampled much earlier in the life cycle. However, the same major microbial families were present within the endosphere of both groups of plants (Figure 2; A, B, C). PCoA plots indicated a shift in the endosphere community when comparing field to pot grown wheat (Figure 1; D1, D2, D3). However, whilst statistical analysis did indicate a significant difference between the overall bacterial and fungal communities associated with the two groups of plants (PERMANOVA, bacterial: R^2^=0.12, *p* < 0.001, fungal: R^2^=0.13, *p* < 0.01, archaeal: R^2^=0.13, *p* > 0.05), subsequent pairwise analysis found no significant difference between any specific compartments (Supplementary Table 2). qPCR indicated that the overall abundance of bacteria and archaea was significantly different between the two groups of plants (*p* < 0.05 in linear models for both microbial groups). While there were significantly more archaea within the bulk soil associated with pot-grown plants (Tukey’s HSD, *p* < 0.01) post-hoc analysis did not show a significant difference in the abundance of either archaea or bacteria in the root associated compartments between the different groups of plants (Tukey HSD, *p* > 0.05 for all). A significantly greater quantity of fungi was detected within the rhizosphere of laboratory-grown plants (Tukey’s HSD, *p* < 0.05) and we also observed lower quantities of all groups within the endosphere (Figure 2; D). Overall, this analysis shows that there is likely a lower microbial abundance within the endosphere of laboratory-grown root growth phase plants, but that any effects on community composition were subtle and mostly restricted to low abundance taxa. As bacterial, fungal, and archaeal communities contained the same major taxa within the endosphere, we conclude that laboratory-grown plants could serve as an approximate experimental analogue for agriculturally cultivated wheat plants when studying the composition of the root microbial community.

### Does *Triticum aestivum* var. Paragon select for specific microbial taxa?

Microbial communities and their functions can differ dramatically between different soils and, as a consequence, soil parameters play a central role in shaping the microbial communities associated with plants [20,48]. To determine if the enrichment of specific microbial taxa and, in particular, the dominance of *Streptomycetaceae* and *Burkholderiaceae*, within the wheat root endosphere was driven by the soil community or by the host, *T. aestivum var.* Paragon was grown in the contrasting soil types (agricultural soil or compost), and a 50:50 mixture of the two. It was reasoned that if *Streptomycetaceae* and *Burkholderiaceae* were dominant only in the agricultural soil and the mixed soil, then certain strains within the agricultural soil might be particularly effective at colonising the endosphere. However, if *Streptomycetaceae* and *Burkholderiaceae* were dominant in the endosphere across all three soil conditions, this would indicate that when present, this family is selectively recruited to the wheat root microbiome. The microbiome was compared between four-week-old (root growth phase) plants grown in Church Farm agricultural soil, Levington F2 compost, and a 50:50 (vol/vol) mix of the two soils under laboratory conditions. Church Farm soil and Levington F2 compost are starkly contrasting soil environments: the agricultural soil is mildly alkaline (pH 7.97), contained only 2.3% organic matter and was relatively low in inorganic nitrogen, magnesium and potassium. Levington F2 compost is acidic (pH 4.98) and has a high organic matter content (91.1%) as well as higher levels of inorganic nitrogen, phosphorus, potassium and magnesium (Supplementary Table 3).

It is well documented that the soil microbial community is a major determinant of endosphere community composition, as endophytic microbes are acquired by plants from the soil [6]. The present study corroborates this observation as PCoA showed clear clustering of communities by soil type, indicating that soil type was an important determinant of the root-associated community composition (Figure 1; E1, E2, E3). For the bacterial and archaeal communities, PERMANOVA corroborated a significant effect of soil type on bacterial community composition for all compartments (Supplementary Table 2). For the fungal community, PERMANOVA also showed significant effect of soil type on the bulk soil and rhizosphere communities (Supplementary Table. 2). For plants cultivated in Levington F2 compost, no data on the fungal community composition within the endosphere could be retrieved. Thus, no statistical comparison could be made. The bacterial communities were distinct between the bulk soil, rhizosphere, and endosphere. This indicated that, while the soil had a significant impact on the composition of the root associated communities, the plant also selects for specific microbial taxa in all the tested soils (Figure 1; E1). PCoA showed a detectable rhizosphere effect (Figure 1; E1) but, consistent with previous studies [24,30], we observed a rhizosphere effect for *T. aestivum* var. Paragon that was subtle as there were only minor differences between the community composition of bulk soil and rhizosphere communities (Figure 2; A, B, C). A SIMPER test revealed that, regardless of soil type, *Streptomycetaceae* (14.6% contribution, *p* < 0.01) and *Burkholderiaceae* (10.1% contribution, *p* < 0.01) were the main taxa driving the community shift from bulk soil to endosphere (Supplementary Table 1). This is supported by the fact that *Streptomycetaceae* and *Burkholderiaceae* were major components of the endosphere bacterial communities under all conditions (Figure 2). Differential abundance analysis demonstrated a significant increase in the abundance of bacterial families *Burkholderiaceae, Chitinophageaceae, Pseudomonadaceae, Rhizobiaceae* and *Streptomycetaceae* within the rhizosphere and/or endosphere across all soil types (Figure 3; C). Enrichment of these groups was correlated with the reduced abundance of some fungal taxa loosely associated with pathogenicity within the endosphere and rhizosphere *(Australiascaceae* [49], *Glomerellaceae* [50,51] and *Hypocreale* [52]), and an increased abundance of one taxon loosely associated with beneficial mycorrhiza *(Leotiaceae* [53–55]) (Figure 3; C).

Further to this, qPCR experiments were performed to compare the abundance of archaea, bacteria, and fungi within the roots of plants cultivated in the agricultural soil or Levington F2 compost. No significant effect of soil type was observed for either fungi or bacteria (ANOVA, *p* > 0.05 for both) (Figure 2; D). However, soil type had a significant effect on the abundance of archaea (*p* < 0.001); there were significantly greater numbers of archaea within the agricultural bulk soil and rhizosphere compartments when compared to those for Levington F2 compost (Tukey’s HSD, *p* < 0.001 for both), but there was no significant difference in the archaeal load detected within the endosphere (Tukey’s HSD, *p* > 0.05). The lower abundance of archaea within Levington F2 compost is surprising given the higher nutrient levels in this soil, and particularly the higher levels of ammonium (Supplementary Table 3).

The archaeal community was dominated by two families of AOA (*Nitrososphaeraceae* and *Nitrosotaleaceae*), which were abundant in all root compartments. *Nitrosotaleaceae* dominated in the more acidic Levington F2 compost whereas *Nitrososphaeraceae* was most abundant in the neutral pH Church Farm soil (Figure 2; C). While soil type was a major determinant of community composition, no selection of specific archaeal lineages within the endosphere was detected by SIMPER or differential abundance analysis, and PCoA did not show a strong effect of compartment on community composition (Figure 1; E2). Contrary to this, there was a small but significant shift in the archaeal community composition overall across compartments (archaeal PERMAN¤VA: R^2^=0.86, *p* = 0.001), and a betadisper analysis was not significant (*p* > 0.01), demonstrating this was not due to difference in dispersion between compartments (Figure 2; C2). Together, these findings might suggest that there is no major selection of archaeal taxa by the wheat roots. However, denaturing gradient gel electrophoresis (DGGE) analysis performed on the archaeal 16S rRNA and *amoA* genes showed a clear shift in the archaeal community across compartments (Supplementary Figure 1). Unfortunately the archaeal 16S rRNA gene database lacks the established framework of its bacterial counterpart [56] and this, coupled with the lack of known diversity or strain characterisation within many archaeal taxa, makes it difficult to achieve good taxonomic resolution from short read amplicon sequencing of the archaeal 16S rRNA gene. We hypothesised therefore that this discrepancy between DGGE and amplicon sequencing arose from the lack of detailed taxonomic representation within the database used to analyse the sequencing data. Despite these limitations, this study has revealed that AOA dominate the archaeal community associated with wheat roots regardless of soil type, and that the abundance of archaea within the root is highest in agricultural soil and increases later in the life cycle of the plant.

### Identification of root exudate utilising microbes using ^13^CO_2_ DNA stable isotope probing

Plants exude 30-40% of the carbon they fix from the atmosphere as root exudates [9]. These compounds can be utilised as a carbon source by microbes residing within and in the vicinity of the root and root exudates could be tailored by the plant to select particular microbial species from the soil. Thus, we aimed to identify the microbial taxa that wheat can support via ^13^CO_2_ DNA SIP. Briefly, wheat was incubated in ^13^CO_2_ for two weeks. During this period, the “heavy” ^13^CO_2_ becomes photosynthetically fixed into carbon-based metabolites and some of these ^13^C labelled compounds are exuded from the roots. Microbial utilisation of these compounds will, in turn, result in the ^13^C label being incorporated into the DNA backbone of actively growing microorganisms. Heavy and light DNA can be separated via density gradient ultracentrifugation and the fractions are then analysed using amplicon sequencing to identify metabolically active microbes. The two-week labelling period was chosen to minimise the probability of labelling via cross feeding by secondary metabolisers [39,57]. Labelling of the bacterial community in the rhizosphere and endosphere was confirmed using DGGE (Supplementary Figures 2 and 5), then heavy and light fractions were pooled and analysed by 16S rRNA gene sequencing (as defined in Supplementary Table 4). The same DGGE experiment was performed using primers targeting archaea and did not indicate labelling and therefore no sequencing of archaea was carried out (Supplementary Figure 3). For fungi, PCR amplification of the ITS2 region for DGGE did not consistently yield products for all fractions, thus DGGE could not be performed. Instead, qPCR was used and did not detect labelling of the fungal community (Supplementary Figure 4) so no sequencing was performed for the fungal community.

PCoA indicated that bacterial communities within endosphere samples were highly variable (Supplementary Figure 5) and there was no significant difference between ^13^C-labelled heavy and light fractions (PERMANOVA: R^2^=0.29, *p* > 0.1). This means the endosphere dataset was too variable to draw any conclusions from the current study about the utilisation of host derived carbon within the endosphere (Supplementary Figure 5). For the rhizosphere however, the replicates were consistent, and PCoA revealed that the bacterial community in the ^13^C heavy fraction was distinct from that of the ^12^C heavy DNA (control) fraction and distinct from the ^13^C DNA and ^12^C light DNA fractions (Figure 4). In addition, the community was significantly different in the ^13^C heavy DNA fraction compared to the unlabelled samples, suggesting that a distinct subset of bacteria was incorporating root-derived carbon (PERMANOVA: R^2^= 0.59, *p* < 0.001). To control for CO_2_ fixation by soil autotrophs the ^13^C heavy fraction was compared to a ^13^C unplanted soil control using PCoA; this analysis indicated that the ^13^C heavy DNA fraction was distinct from the ^13^C bulk soil control (Figure 4). After these comparisons, we could be confident that the shift in community composition within the ^13^C heavy DNA fraction was driven by microbes within the rhizosphere actively utilising root exudates. Differential abundance analysis was performed to identify the taxa driving these shifts. Exudate metabolisers were defined as taxa showing significantly greater abundance within ^13^C heavy DNA fractions when compared with both the ^13^C light fractions and the ^12^C control heavy fractions. Above the abundance threshold, we identified 9 exudate-utilising bacterial taxa (Figure 6, Supplementary Table 5). While *Streptomycetaceae* were not among these, *Pseudomonadaceae* were utilising root exudates, as were two other bacterial taxa *(Comamonadaceae* and *Oxalobacteriaceae*) which likely belonged to the *Burkholderiaceae*. As defined by the Genome Taxonomy Database [58], *Comamonadaceae* and *Oxalobacteriaceae* are now classified as genera *Comamonas* and *Oxalobacter* within the *Burkholderiaceae* family. Within the ^13^C unplanted soil control, differential abundance analysis indicated that six taxa were significantly enriched in the heavy DNA fraction compared to the light fraction these taxa are hypothesised to fix ^13^CO_2_ autotrophically (Supplementary Table 10). Only one taxon was ^13^C-labelled in both the rhizosphere and unplanted soil, *Intrasporangiaceae,* and thus was excluded from the list of root exudate utilising bacterial taxa. While microbes belonging to this family are capable of photosynthesis, they also have genomes with high GC content, and as such they may be overrepresented in heavy fractions.

**Figure 4.**
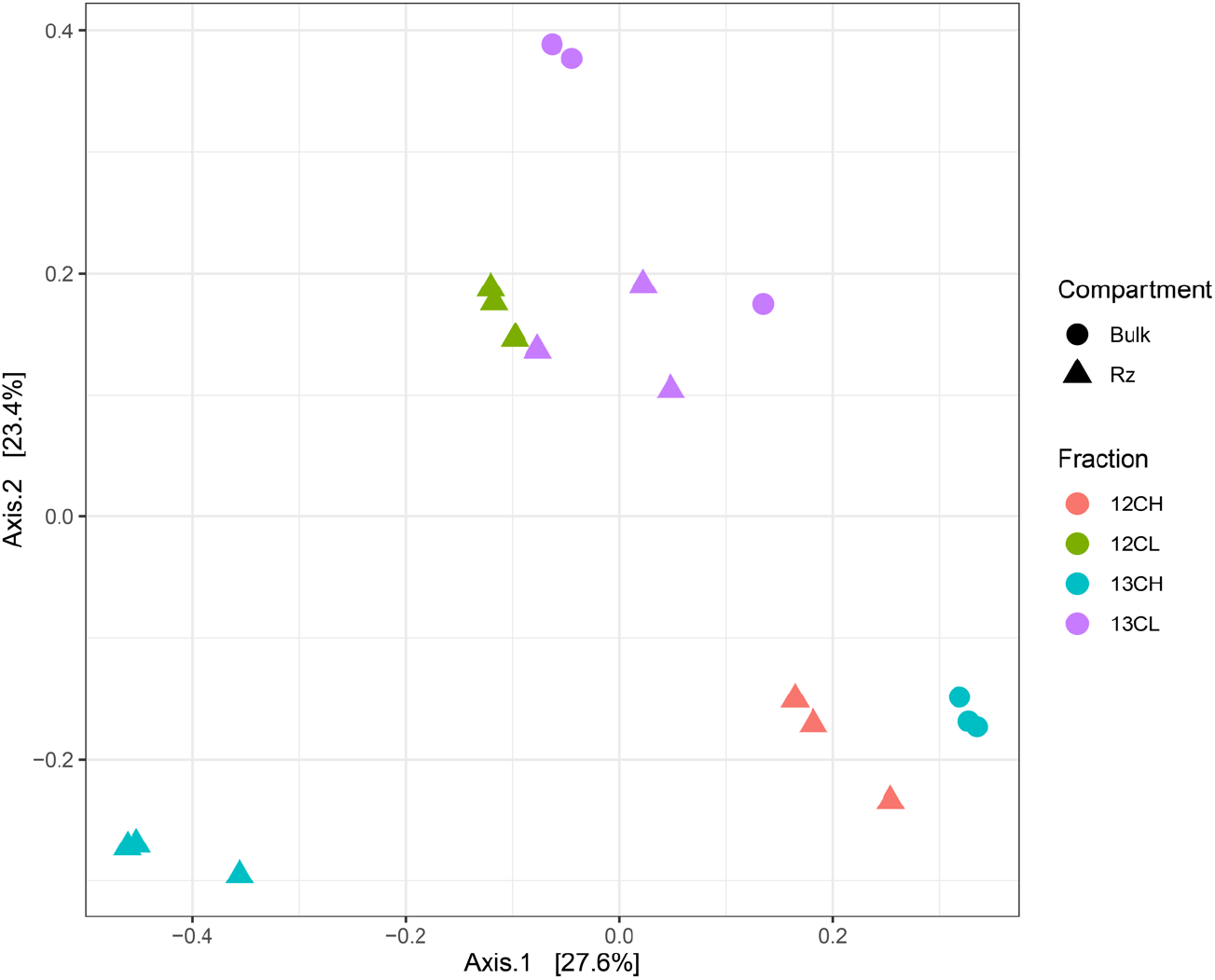
Principle coordinates analysis (PCoA) of Bray-Curtis dissimilarity between bacterial families present in the heavy and light fractions of rhizosphere and bulk soil ^13^C labelled and ^12^C unlabelled treatments (N=3 replicate plants per CO_2_ treatment). Rhizosphere communities were shown to vary significantly between labelled or unlabelled fractions (PERMANOVA: permutations=999, R^2^= 0.59, *p* < 0.001).

**Figure 5.**
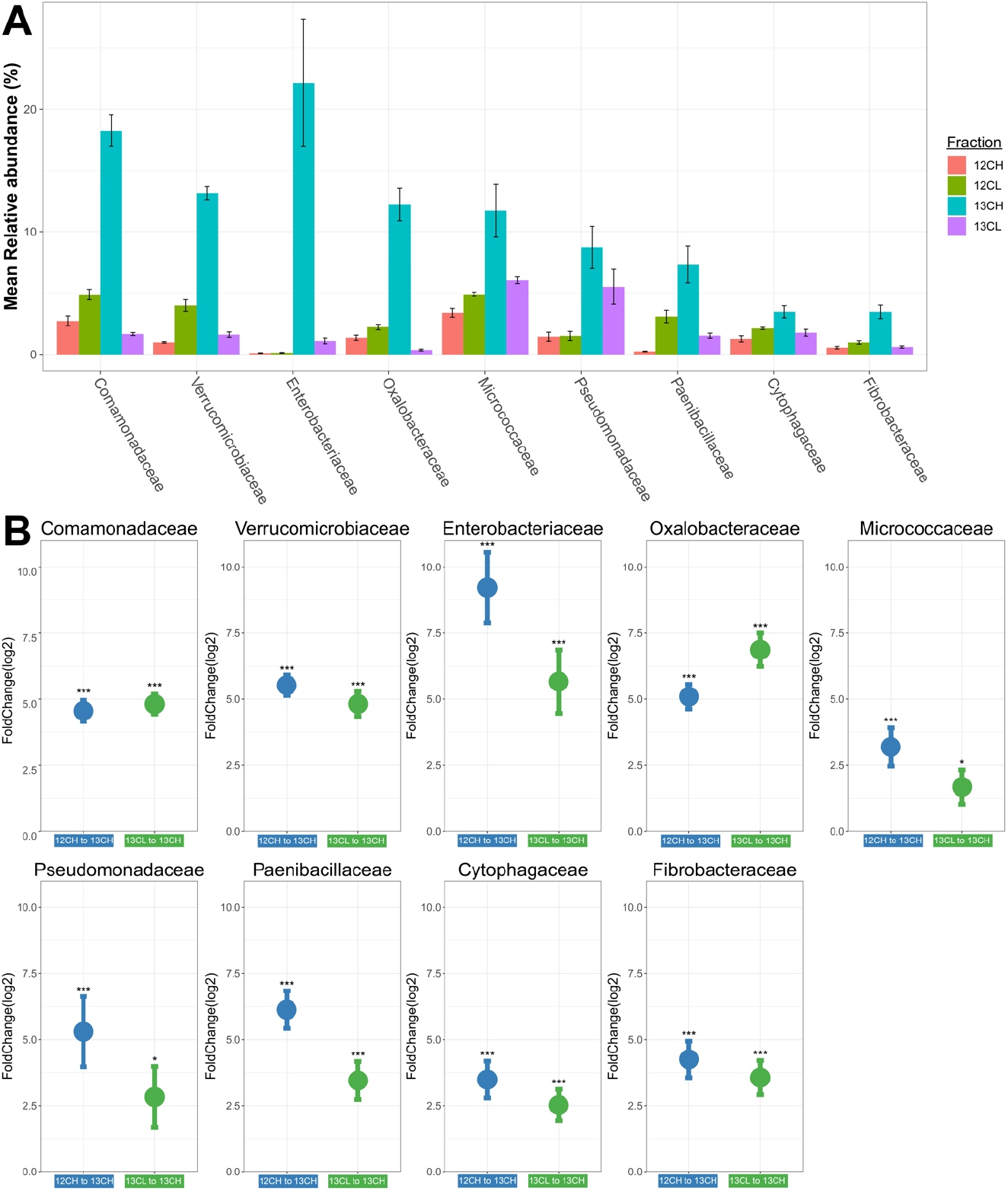
**A** Mean relative abundance of each bacterial family in the rhizosphere of plants incubated with ^12^CO_2_ or ^13^CO_2_. N=3 replicate plants per treatment. Bars represent ± standard errors of the mean. **B** The results of differential abundance analysis for bacterial families in the rhizosphere; points show the log2 fold change of different bacterial families between the ^12^CO_2_ heavy and the ^13^CO_2_ heavy fraction (blue) or between the ^13^CO_2_ light and the ^13^CO_2_ heavy fraction (green) (N=3). Log2-fold change standard errors of triplicate plants is shown. *** represents taxa with a significant log2fold change *(p* < 0.001) For the full statistical output see Supplementary Table 5.

## Discussion

In this work we profiled the microbial communities in the rhizosphere and endosphere of the UK elite Spring bread wheat *T. aestivum* variety Paragon. We identified the core microbial families associated with the rhizosphere and endosphere of these plants and the subset of microorganisms assimilating plant-derived carbon in the rhizosphere. This study revealed that plant developmental senescence induces shift in the root-associated microbial communities and an increase in microbial abundance in the plant endosphere. Concurrent with established literature [6,21,59] we found the soil inoculum to be a major driver of root community composition. Given the contrasting range of soils, wheat varieties, developmental timepoints, and growth management strategies used across studies, drawing direct comparisons is often challenging. For example Schlatter *et al.* identified *Oxolabacteraceae, Comamonadaceae* and *Chitinophaga* as core rhizobacteria for the wheat cultivar *Triticum aestivum* L. cv. Louise [29]. Our work corroborates this observation for *T.* aestivum *var*. Paragon, all these taxa were identified by SIP as exudate utilising microbes. However, many of the core taxa identified by Schlatter *et al.* were not identified by the present work. Similarly, for the endosphere community, Kuźniar and colleagues identified *Flavobacterium, Janthinobacterium,* and *Pseudomonas* as core microbiota for both cultivars tested, and *Paenibacillus* as a core taxon for *T. aestivum* L. cv. Hondia [28]. We identified *Pseudomonadaceae* as a core component of the *T. aestivum* var. Paragon endosphere microbiome and, while *Paenibacillaceae* were not enriched in the endosphere consistently, we did identify this family as an exudate utiliser within the rhizosphere. *Streptomycetaceae* were not identified by the study of Kuźniar and colleagues. While these combined results consistently imply a role for common taxa such as *Pseudomonadaceae* or members of the *Burkholderiaceae* family, it cannot explain the differences observed in colonisation by other taxa, and in particular *Streptomycetaceae.* While it is likely this is largely driven by soil type, there is some evidence that for wheat, similarly to barley [11], plant genotype may be responsible for these differences [24,28,32,44]. In a study which used the same Church Farm field site as our work, *T. aestivum var.* Paragon was previously reported to be an outlier compared to other wheat varieties, with a particularly distinct rhizosphere and endosphere community [24]. Further studies are needed to fully assess how wheat rhizosphere and endosphere communities vary across different wheat cultivars and soil environments, and which of these factors has the greatest influence.

While only slight differences were observed between root-growth phase laboratory cultivated plants and stem elongation phase field cultivated plants, significant changes in the abundance of numerous bacterial and fungal taxa occurred at the onset of plant developmental senescence. To our knowledge, the wheat root community has not previously been assessed after senescence, though development has been shown to significantly alter the wheat rhizosphere community [22,23]. One fungal group, *Chaetosphaeriaceae,* was significantly enriched as the plant senesced. This family represents a relatively diverse group of fungi, although members of this group such as *Chaetosphaeria* are known to reproduce within decomposing plant tissues, which may explain the four-fold increase in abundance after senescence [60]. In terms of the overall fungal community composition (Figure 2; B1), the greatest change during senescence was in the Pleosporales group, and this may also contribute to the observed increase in fungal abundance during senescence. This group was excluded from the differential abundance analysis which focused on lower taxonomic ranks. Pleosporales is an order of fungi containing over 28 families [61], and such a high diversity makes the ecological role of this group difficult to postulate. Some families within the Pleosporales are associated with endophytic plant parasites [61], including necrotrophic pathogens of wheat *Pyrenophora tritici-repentis* and *Parastagonospora nodorum* [62]. Necrotrophic pathogens specialise in colonising and degrading dead plant cells, and senescent tissues are thought to provide a favourable environment for necrotrophs [37]. It is interesting to note that this increased fungal colonisation correlated with reduced abundance of fungi-suppressive endophytic bacteria such as *Streptomycetaceae* [63,64] and *Burkholderiaceae* [65] during developmental senescence. The present work, however, cannot provide any direct evidence of a causative relationship driving this correlation.

*Burkholderiaceae*-family taxa (*Comamonadaceae and Oxalobacteriaceae*), and *Pseudomonadaceae* were identified as potential root exudate utilisers within the rhizosphere, in agreement with previous studies [42,66]. These bacterial groups were also consistently enriched in the rhizosphere or endosphere, regardless of soil type. These results imply these families may be selectively recruited to the plants via root exudates, which support *Burkholderiaceae* and *Pseudomonadaceae* via photosynthetically fixed carbon. The *Pseudomonadaceae* family contains a diverse range of plant-beneficial and plant pathogenic strains [67,68] but the literature correlates exudate utilisation with microbial functions which benefit the host plant [69,70], and exudates can have a negative effect on plant pathogens [12]. While the mechanism of this selectivity remains unknown, it is likely these exudate utilisers are plant beneficial strains. Well studied representatives of this family with plant growth promoting traits include *Pseudomonas brassicacearum* [71] and *Pseudomonas fluorescens* [72]. Most of the exudate utilising families identified in the present work were fast growing Gram-negative bacteria. As observed by Worsley and colleagues (bioRxiv [73]), faster growing organisms are labelled more readily within a two-week incubation period. Due to their faster growth rates, these microorganisms can more easily monopolise the plant derived carbon within the rhizosphere and incorporate ^13^C into the DNA backbone during DNA replication. Slower growing organisms such as *Streptomycetaceae* are likely outcompeted for root derived resources in the rhizosphere or the two-week incubation period may be too short to allow the incorporation of the ^13^C label into DNA.

*Streptomycetaceae* were the most abundant of the core endosphere enriched families, despite not incorporating root derived carbon in the rhizosphere. This family is dominated by a single genus, *Streptomyces.* These filamentous Gram-positive bacteria are well known producers of antifungal and antibacterial secondary metabolites, and members of the genus have been shown to promote plant growth [64], have been correlated with increased drought tolerance [74], and can protect host plants from disease [63,64]. *Streptomyces* species make up the active ingredients of horticultural products Actinovate and Mycostop and it has been proposed that plant roots may provide a major niche for these bacteria which are usually described as free-living, soil dwelling saprophytes. In this study *Streptomycetaceae* accounted for up to 40% of the bacteria present in the endosphere for some plants. Intriguingly, after the plants senesced, there was a two-fold reduction in the abundance of *Streptomycetaceae* within the endosphere. This a surprising result for a bacterial group typically associated with the breakdown of dead organic matter within soils [75]. As plants senesce and die, a process of ecological succession occurs, where the tissues are colonised by different microbes (particularly fungi) successively as different resources within the plant tissues are degraded [76,77]. The first microorganisms to colonise will be those rapidly metabolising sugars and lipids, followed later by more specialist organisms which will breakdown complex molecules like lignin and cellulose. While these later stages are typically attributed to fungi, *Streptomycetaceae* are known to degrade complex plant derived molecules such as hemicellulose and insoluble lignin [75,78]. It could be that our sampling timepoint (late in the developmental senescence process, but prior to most biomass degradation) was too early in this succession process for any biomass fuelled *Streptomycetaceae* proliferation to be obvious. This however cannot explain the reduced abundance of *Streptomycetaceae* in senesced roots compared to the actively growing plants. This might be explained by a lack of active input from the plant, as the host senesces and resources are diverted to the developing grain [35] host derived resources may no longer be available to support *Streptomycetaceae* growth in the endosphere. The DNA-SIP experiment indicated that *Streptomycetaceae* did not utilise root exudates under the selected experimental conditions, which contradicts the findings of Ai and colleagues [43]. It must be noted that while *Streptomycetaceae* were not labelled in the DNA-SIP experiment, this experiment focused on the rhizosphere, and our data demonstrated that *Streptomycetaceae* primarily colonise the endosphere.

Further SIP experiments exploring the endosphere community, with more replicates to account for the high variability, may help to determine whether *Streptomycetaceae* can utilise plant derived carbon within the endosphere, and if the loss of these resources explains their reduced presence during senescence. Future studies should also investigate how *Streptomycetaceae* are able to colonise and survive within the endosphere of wheat. During developmental senescence, nitrogen is the main resource diverted to the developing grain [35]. It is possible that nitrogen, not carbon, is the resource provided by the host plant to support *Streptomycetaceae* growth. There is precedent for host-derived metabolites such as amino acids or gamma-aminobutyric acid (GABA) acting as a nitrogen source for root associated microbes [70,79]. Additionally, there is evidence that the increased use of nitrogen fertilizer (which correlates with greater total root exudation) was negatively correlated with the abundance of *Streptomycetaceae* in the rhizosphere [23]. In the future, ^15^N-nitrogen DNA or RNA-SIP could be used to explore whether *T. aestivum var.* Paragon is able to support *Streptomycetaceae* within the endosphere via nitrogen containing, host-derived metabolites.

In conclusion: (1) We identified five core microbial taxa associated within the rhizosphere and endosphere of *T. aestivum var.* Paragon, *Streptomycetaceae, Burkholderiaceae, Pseudomonadaceae, Rhizobiaceae* and *Chitinophageaceae.* The consistency of the enrichment of these groups across the soil types and plant growth stages we tested strongly indicates that they are core taxa associated with Paragon *var. T. aestivum*. This, however, cannot be extrapolated to other varieties of wheat, and one study even suggests *T. aestivum* var. Paragon is an outlier with a particularly distinct microbiome [24]. To gain a more detailed understanding of which microbial taxa are associated with the roots of spring bread wheat, more genotypes must be analysed. (2) At the onset of developmental senescence, significant reductions in the abundance of many taxa were observed, including the whole core endosphere and rhizosphere microbiome. In particular, *Streptomycetaceae* abundance was reduced two-fold. This may indicate that active input from the host is required to maintain the abundance of certain families within the endosphere. A significant increase in the total abundance of bacteria and archaea was evident during senescence and potentially increased colonisation of fungal groups associated with necrotrophy and plant tissue degradation. (3) No lineages of archaea were specifically associated with wheat roots. Conflicting data from DGGE and from 16S rRNA gene sequencing indicated that the currently available archaeal 16S rRNA gene databases are not sufficiently complete for this metabarcoding approach. In the future, longer read methods or metagenomics could be applied to better investigate archaeal community dynamics within the root microbiome. (4) We identified nine taxa within the rhizosphere utilising carbon from wheat root exudates, including aforementioned core taxa of *T. aestivum* var. Paragon, *Pseudomonadaceae* and *Burkholderiaceae.* There was no evidence that the most abundant endosphere bacterial family *Streptomycetaceae* was using plant exudates within the rhizosphere. Future ^13^CO_2_ SIP experiments should utilise a higher number of replicates to account for endosphere variation and to identify which families utilise host-derived carbon inside the root. Given the reduction in *Streptomycetaceae* abundance during senescence, future work should also consider exploring host derived nitrogen as a potential medium through which *T. aestivum* var. Paragon might support endophytic bacteria. The present work has provided novel insights into the composition and variation within the wheat microbiome and how the community changes through developmental senescence. Greater understanding is needed of the role played by the five core taxa associated with *T. aestivum var.* Paragon, and the mechanisms by which they are able to colonise the root and are supported by the host. This knowledge may inform novel agricultural applications or more ecologically responsible management strategies for wheat.

## Methods

### Soil sampling and chemical analyses

Agricultural soil was sampled in April 2019 from the John Innes Centre (JIC) Church Farm cereal crop research station in Bawburgh (Norfolk, United Kingdom) (52°37’39.4”N 1°10’42.2”E). The top 20cm of soil was removed prior to sampling. Levington F2 compost was obtained from the John Innes Centre. Soil was stored at 4°C and pre-homogenised prior to use. Chemical analysis was performed by the James Hutton Institute Soil Analysis Service (Aberdeen, UK) to measure soil pH, organic matter (%), and the phosphorus, potassium, and magnesium content (mg/kg) (Supplementary Table 3). To quantify inorganic nitrate and ammonium concentrations a KCl extraction was performed where 3g of each soil type suspended in 24ml of 1 M KCl in triplicate and incubated for 30 minutes with shaking at 250rpm. To quantify ammonium concentration (g/kg) the colorimetric indophenol blue method was used [80]. For nitrate concentration (g/kg) vanadium (III) chloride reduction coupled to the colorimetric Griess reaction as previously described in Miranda *et al.* [81].

### Wheat cultivation, sampling and DNA extraction

Paragon var. *Triticum aestivum* seeds were soaked for two minutes in 70% ethanol (v/v), 10 minutes in 3% sodium hypochlorite (v/v) and washed 10 times with sterile water to sterilise the seed surface. Seeds were sown into pots of pre-homogenised Church farm agricultural soil, Levington F2 compost, or a 50:50 (v/v) mix of the two. Plants were propagated for 30 days at 21°C under a 12 h light/ 12 h dark photoperiod before endosphere, rhizosphere and bulk soil samples were analysed. To assess microbial community diversity in the field, Paragon var. *Triticum aestivum* plants were sampled during the stem elongation growth phase approximately 200 days after sowing, in July 2019. To assess microbial diversity after senescence, three Paragon var. *Triticum aestivum* plants were sampled immediately before harvest in August 2020 approximately 230 days after sowing. All field grown plants were sampled from the JIC Church Farm field studies site in Bawburgh (Norfolk, United Kingdom) (52°37’42.0”N 1°10’36.3”E) and were cultivated in the same field from which agricultural soil was sampled.

Microbial communities were analysed in the bulk soil, rhizosphere and endosphere for all plants. All three compartments were analysed from triplicate plants for each condition described (Church farm agricultural soil, Levington F2 compost, 50:50 vol/vol mix, and field-grown stem elongation or senescence). After a plant was removed, the potted soil associated with each plant was homogenised and a bulk soil sample was taken. For field-grown wheat bulk soil samples were taken from unplanted soil approximately 30cm away from the plant, in the same way as described for soil sampling. For all plants the phyllosphere was removed using a sterile scalpel and discarded. To analyse the rhizosphere and endosphere samples, loose soil was lightly shaken off of the roots, then roots were washed in phosphate buffer saline (PBS) (6.33g NaH_2_PO_4_.H_2_O, 16.5g Na_2_HPO_4_.H_2_O, 1L dH_2_O, 0.02% Silwett L-77 (v/v)). Pelleted material from this wash was analysed as the rhizosphere sample. To obtain the endosphere samples, remaining soil particles were washed off of the roots with PBS buffer. Then roots were soaked for 30 seconds in 70% ethanol (v/v), 5 minutes in 3% sodium hypochlorite (v/v) and washed 10 times with sterile water for surface sterilisation. To remove the rhizoplane roots were then sonicated for 20 minutes in a sonicating water bath [6]. After processing, all root, rhizosphere, and soil samples were snap frozen and stored at −80°C. The frozen root material was ground up in liquid nitrogen with a pestle and mortar. For all samples DNA was extracted using the FastDNA™ SPIN Kit for Soil (MP Biomedical) according to manufacturer’s protocol with minor modifications: incubation in DNA matrix buffer was performed for 12 minutes and elution carried out using 75μl DNase/Pyrogen-Free Water. All DNA samples were stored at −20°C. DNA quality and yields were assessed using a nanodrop and Qubit fluorimeter.

### ^13^C CO_2_ labelling of wheat for DNA SIP

Agricultural soil was sampled in July 2019, sampling method was as previously described. The soil was homogenized; any organic matter, or stones larger than ~3cm, were removed before soil was spread out to a depth of ~2cm and dried at 20°C overnight. Soil was added to pots and wetted before surface sterilized *T. aestivum var.* Paragon seeds were sown (surface sterilisation performed as described), three additional pots remained unplanted as controls for autotrophic CO_2_ fixation by soil microorganisms. Plants were grown in unsealed gas tight 4.25L PVC chambers under a 12 h light/ 12 h dark photoperiod at 21°C for 3 weeks. Then at the start of each photoperiod the chambers were purged with CO_2_ free air (80% nitrogen, 20% oxygen, British Oxygen Company, Guilford, UK) and sealed before pulse CO_2_ injection every hour. During each photoperiod 3 plants and 3 unplanted soil controls were injected with ^13^C CO_2_ (99% Cambridge isotopes, Massachusetts, USA) and 3 plants were injected with ^12^C CO_2_. Headspace CO_2_ was maintained at 800ppmv (~twice atmospheric CO_2_). Plant CO_2_ uptake rates were determined every 4 days to ensure the volume of CO_2_ added at each 1 h interval would maintain approximately 800ppmv. For this, headspace CO_2_ concentrations were measured using gas chromatography every hour. Measurements were conducted using an Agilent 7890A gas chromatography instrument, with flame ionization detector, a Poropak Q (6ft x 1/8”) HP plotQ column (30m x 0.530mm, 40μm film), a nickel catalyst, and a helium carrier gas. The instrument ran with the following settings: injector temperature 250°C, detector temperature 300°C, column temperature 115°C and oven temperature 50°C. The injection volume was 100μl and run time was 5mins (CO_2_ retention time is 3.4 mins). A standard curve was used to calculate CO_2_ ppmv from peak areas. Standards of known CO_2_ concentration were prepared in nitrogen flushed 120ml serum vials. The volume of CO_2_ injected at each 1h interval to maintain 800ppmv CO_2_ was calculated as follows: Vol CO_2_ (ml) = (800 (ppmv) – headspace CO_2_ after 1 hour (ppmv) / 1000) * 4.25(L). At the end of each photoperiod, tube lids were removed to prevent build-up of CO_2_ during the dark period. At the start of the next 12 h, photoperiod tubes were flushed with CO_2_ free air and headspace CO_2_ was maintained at 800ppmv as described. After 14 days of labelling for all plants bulk soil, rhizosphere, and endosphere compartments were sampled as described previously and snap-frozen prior to DNA extraction as described previously.

### Density gradient ultracentrifugation and fractionation for DNA SIP

Density gradient ultracentrifugation was used to separate ^13^C labelled DNA from ^12^C DNA as previously described by Neufeld and colleagues [57]. Briefly, for each sample 700ng of DNA was mixed with a 7.163 M CsCl solution and gradient buffer (0.1M Tris-HCl pH8, 0.1M KCl,1mM EDTA) to a final measured buoyant density of 1.725 g/ml^-1^. Buoyant density was determined via the refractive index using a refractometer (Reichert Analytical Instruments, NY, USA). Samples were loaded into polyallomer quick seal centrifuge tubes (Beckman Coulter) and heat-sealed. Tubes were placed into a Vti 65.2 rotor (Beckman-Coulter) and centrifuged for 62 hours at 44,100 rpm (~177,000gav) and 20°C under a vacuum. Samples were fractionated by piercing the bottom of the ultracentrifuge tube with a 0.6 mm sterile needle and dH2O was pumped into the centrifuge tube at a rate of 450μl per minute, displacing the gradient into 1.5ml microcentrifuge tubes. Fractions were collected until the water had fully displaced the gradient solution; this resulted in twelve 450μl fractions. The DNA was precipitated from fractions by adding 4μl of Co-precipitant Pink Linear Polyacrylamide (Bioline) and 2 volumes of PEG-NaCl solution (30% w/v polyethylene glycol 6000, 1.6M NaCl) to each fraction, followed by an overnight incubation at 4°C. Fractions were then centrifuged at 21,130g for 30 minutes and the supernatant was discarded. The DNA pellet was washed in 500μl 70% EtOH and centrifuged at 21,130 g for 10 minutes. The resulting pellet was air-dried and resuspended in 30μl sterile dH2O. Fractions were then stored at −20°C. Fractions were pooled prior to sequencing (supplementary table 4), sequencing was performed as described in the DNA sequencing and analysis section, except that peptide nucleic acid (PNA) blockers were used to prevent amplification of chloroplast and mitochondrial 16S rRNA genes.

### DNA sequencing and analysis

All 16S rRNA genes were amplified using primers specific to the archaeal (A0109F/A1000R) or bacterial (PRK341F/MPRK806R) gene (Supplementary Table 6). The fungal 18S ITS2 region was amplified using primers specifically targeting fungi (fITS7Fw/ITS4Rev_2) to avoid *Triticum aestivum* ITS2 amplification (Supplementary Table 6). No fungal ITS2 amplicon could be obtained from the endosphere of Levington F2 compost plants. PCR conditions are indicated in Supplementary Table 7. Purified PCR products were sent for paired-end sequencing using an Illumina MiSeq platform at Mr DNA (Molecular Research LP, Shallowater, Texas, USA). The bacterial 16S rRNA gene was sequenced using the PRK341F/MPRK806R primers (465bp). The archaeal 16S rRNA gene was sequenced using the A0349F/A0519R primers (170bp). The fungal ITS2 region was sequenced with the fITS7Fw/ITS4Rev_2 primers (350bp). See Supplementary Table 6 for primer sequences. Upon receipt, all sequencing reads were further processed using the software package quantitative insights into microbial ecology 2 (Qiime2 [82]) version 2019.7. Paired-end sequencing reads were demultiplexed and then quality filtered and denoised using the DADA2 plugin version 1.14 [83]. Reads were trimmed to remove the first 17-20 base pairs (primer dependent, see Supplementary Table 8) and truncated to 150-230 base pairs to remove low quality base calls (dependent on read quality and amplicon length, see Supplementary Table 8). Chimeras were removed using the consensus method and default settings were used for all other analyses. For taxonomic assignments bayesian bacterial and archaeal 16S sequence classifiers were trained against the SILVA [84] database version 128 using a 97% similarity cut off. For the fungal ITS2 reads, the bayesian sequence classifier was trained against the UNITE [85] database version 8.0 using a 97% similarity cut-off. Taxonomy-based filtering was performed to remove contaminating mitochondrial, chloroplast and *Triticum* sequences (Supplementary Table 9), remaining sequences were used for all further analyses. Taxonomy-based filtering was not required for the fungal dataset.

Statistical analysis was performed using *R* version 3.6.2 [86]. The package vegan version 2.5-7 [87] was used to calculate Bray Curtis dissimilarities and conduct similarity percentages breakdown analysis (SIMPER [88]). Permutational Multivariate Analysis of Variance (PERMANOVA) analyses were conducted using Bray Curtis dissimilarity matrices and the *adonis* function in vegan. Bray Curtis dissimilarities were also used for principle coordinate analysis (PCoA) which was performed using the packages phyloseq version 1.3 [89] and plyr. Differential abundance analysis was performed using DESeq2 in the package microbiomeSeq version 0.1 [90]. Given the low number of reads which remained in some samples after taxonomy-based filtering (Table 9), a base mean cut off of 200 for the field and pot metabarcoding experiments, or of 400 for the stable isotope probing experiment, was applied to the DESeq2 output to eliminate possible false positives resulting from low sequencing depth. If a taxon had a base mean > 200 and a significant p-value in one or more comparison, data for that taxon was plotted in Figure 3 for all comparisons. For details see Supplementary Tables 4, 10-16.

### Real-time quantitative PCR

The abundance of bacterial or archaeal 16S rRNA genes and of fungal 18S rRNA genes was determined by qPCR amplification of these genes from DNA extracts. Bacterial 16S rRNA abundance was quantified using bacteria-specific primers Com1F/769r, as previously described [91]. Archaeal 16S rRNA gene abundance was quantified using the archaeal specific A771f/A957r primers, as previously described [92]. Fungi-specific primers, as previously described [93], FR1F/FF390R were used to quantify 18S rRNA gene abundance and examine ^13^C labelling of the fungal community for the SIP fractions. Primer sequences are presented in Supplementary Table 6. The qPCR was performed using the Applied Biosystems QuantStudio 1 Real-Time PCR System (Applied Biosystems, Warrington, UK) with the New England Biolabs SYBR Green Luna^®^ Universal qPCR Master Mix (New England Biolabs, Hitchin, UK). PCR mixtures and cycling conditions are described in Supplementary Table 7. Bacterial, fungal and archaeal qPCR standards were generated using a set of primers enabling amplification of the full length bacterial or archaeal 16S rRNA gene or fungal 18S rRNA gene, cloned into the Promega pGEM^®^-T Easy Vector system, and the correct sequence was validated by Sanger sequencing (Supplementary Table 6). After purification, the standard was diluted from 2×10^7^ to 2×10^0^ copies/μl in duplicate and ran alongside all qPCR assays. Ct values from standard dilutions were plotted as a standard curve and used to calculate 16S/18S rRNA gene copies/50 ng DNA extract. Amplification efficiencies ranged from 90.9% to 107% with R^2^ > 0.98 for all standard curve regressions. All test samples were normalised to 50ng of template DNA per reaction and ran in biological triplicate. PCR products were all analysed by both melt curves and agarose gel electrophoresis which confirmed amplification of only one product of the expected size. For statistical comparison of the average 16S rRNA or 18S rRNA gene copy number between samples ANOVA and linear models, followed by Tukey post-hoc was run in *R* [86].

### Denaturing gradient gel electrophoresis (DGGE)

DGGE was performed separately on the bacterial and archaeal 16S rRNA genes to screen SIP fractions for a change in the community in the heavy compared to the light fractions, and between the ^13^CO_2_ labelled heavy fractions and those of the ^12^CO_2_ control plants. A nested PCR approach was taken to amplify the archaeal 16S rRNA gene, the first round used primers A109F/A1000R and the second introduced a 5’ GC clamp using A771F-GC/A975R (Supplementary Table 6). The same method was used to screen for a shift in the archaeal community across root compartments. One round of PCR was used for bacterial DGGE using the primers PRK341F-GC/518R to introduce a 5’ GC clamp, and for archaeal *amoA* DGGE using CrenamoA23f/A616r (Supplementary Table 6). PCR conditions are indicated in Supplementary Table 7. An 8% polyacrylamide gel was made with a denaturing gradient of 40-80% (2.8M urea / 16% (vol/vol) formamide, to 5.6M urea / 32% (vol/vol) formamide), and a 6% acrylamide stacking gel with 0% denaturant. 2-8μl of PCR product was loaded per well for each sample and the gel was loaded into an electrophoresis tank filled with 1x Tris acetate EDTA (TAE) buffer (242g Tris base, 57.1ml acetic acid, 100ml 0.5M EDTA pH 8.0). Electrophoresis ran at 0.2 amps, 75 volts and 60°C for 16 hours. After washing, gels were stained in the dark using 4μl of SYBR gold nucleic acid gel stain (Invitrogen™) in 400ml 1x TAE buffer. After one hour, gels were washed twice before imaging using a Bio-Rad Gel Doc XR imager.

## Declarations

### Ethics approval and consent to participate

Not Applicable.

### Consent for publication

Not Applicable.

### Availability of data and material

The datasets generated during and/or analysed during the current study are available in the European Nucleotide Archive. Accession number PRJEB42686 (https://www.ebi.ac.uk/ena/browser/view/PRJEB42686).

### Authors’ contributions

SMMP, JCM, LLM and MIH designed the metabarcoding experiments and all authors contributed to the design of the stable isotope probing (SIP) experiment. SMMP performed all the metabarcoding and qPCR experiments, and SMMP and SFW performed all subsequent bioinformatic and statistical analysis. SP assessed the fungal and archaeal communities for the SIP experiment. SMMP and JTN performed the labelling, and the density gradient ultracentrifugation and fractionation for the SIP experiment. JTN performed bacterial community denaturing gradient gel electrophoresis, metabarcoding, and all subsequent bioinformatic and statistical analysis for the SIP experiment. Field sampling was performed by SMMP, JTN, and MIH. All authors contributed to the development of and approved the final manuscript.

## Acknowledgements

S.M.M.P. and S.F.W. were funded by Natural Environment Research Council (NERC) PhD studentships (NERC Doctoral Training Programme grant NE/L002582/1). J.T.N. was funded by a Biotechnology and Biological Sciences Research Council (BBSRC) PhD studentship (BBSRC Doctoral Training Program grant BB/M011216/1). L.L.M is supported by a Royal Society Dorothy Hodgkin Research Fellowship (DH150187) and by a European Research Council (ERC) Starting Grant (UNITY 852993). DNA sequencing was performed at Molecular Research Ltd and we thank Dr Scot E. Dowd for this service. Seed materials were acquired from the John Innes Centre Germplasm Resource Unit (JIC GRU) and we thank the whole GRU team for their invaluable support with this work. Field studies were performed at the John Innes Centre Field Studies Site in Bawburgh, and we thank the Simon Orford for invaluable help with field sampling, and the entire John Innes Centre Cereal Crop Research team for all their support. Computational analysis was performed using the High-Performance Computing Cluster supported by the Research and Specialist Computing Support service at the University of East Anglia and we would like to thank the entire team for their support with this work. We would like to thank Dr Richard Oliver for valuable conversations on fungal taxonomy and fungal community dynamics during senescence.

