## Supplementary information for "Soil, senescence and exudate utilisation: Characterisation of the Paragon var. spring bread wheat root microbiome"

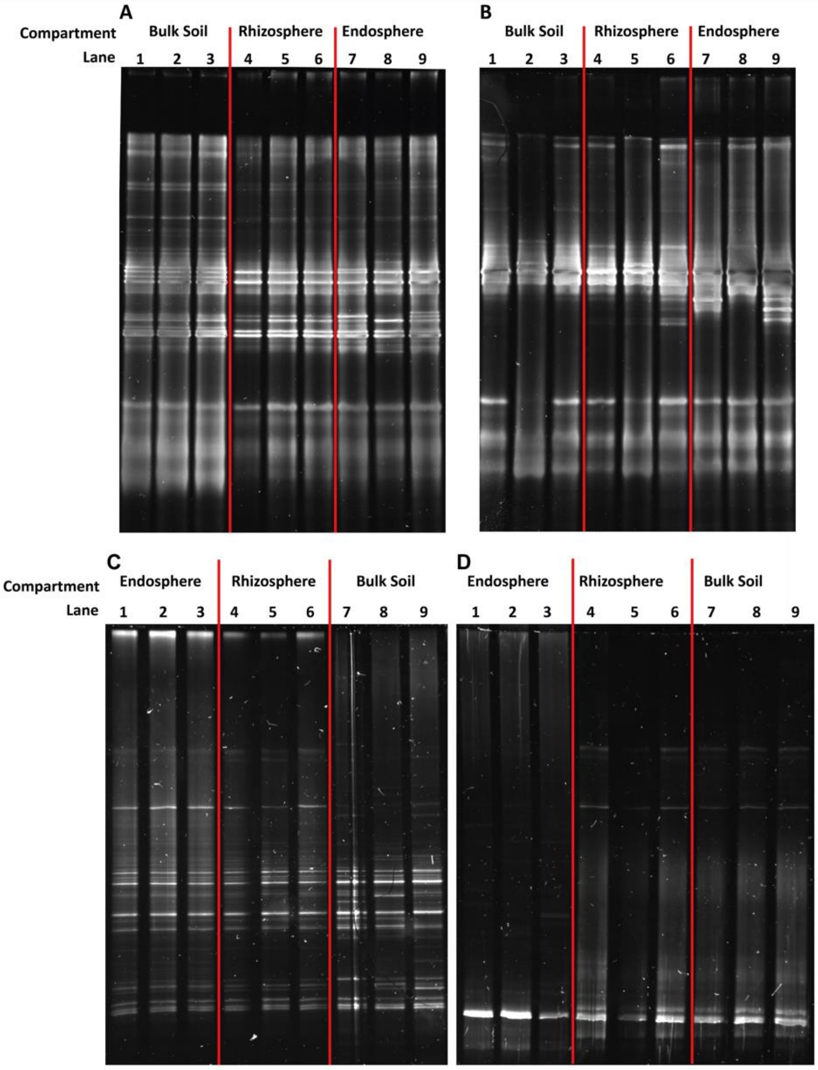


**Supplementary Figure 1.** Denaturing gel gradient electrophoresis (DGGE) showing archaeal 16S rRNA gene (A, B) or *amoA* (C, D) diversity across the bulk soil, rhizosphere and endosphere of wheat grown under laboratory conditions in agricultural soil (A, C) or Levington F2 compost (B, D).Primers are indicated in the Supplementary Table 6.

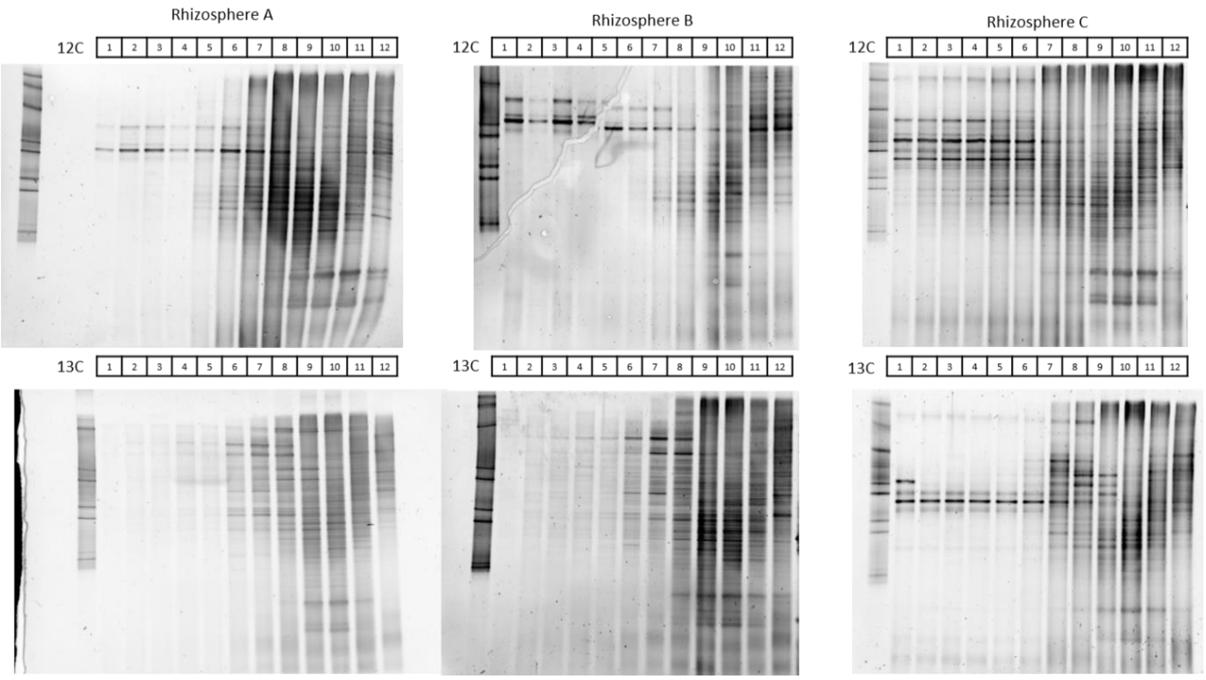


**Supplementary Figure 2**. Denaturing gel gradient electrophoresis (DGGE) showing bacterial 16S rRNA gene diversity across the 12 fractions generated for stable isotope probing for the rhizosphere associated with the $^{12}C$ control (top) and $^{13}C$ labelled (top) plant (n=3). Primers indicated in Supplementary Table 6.

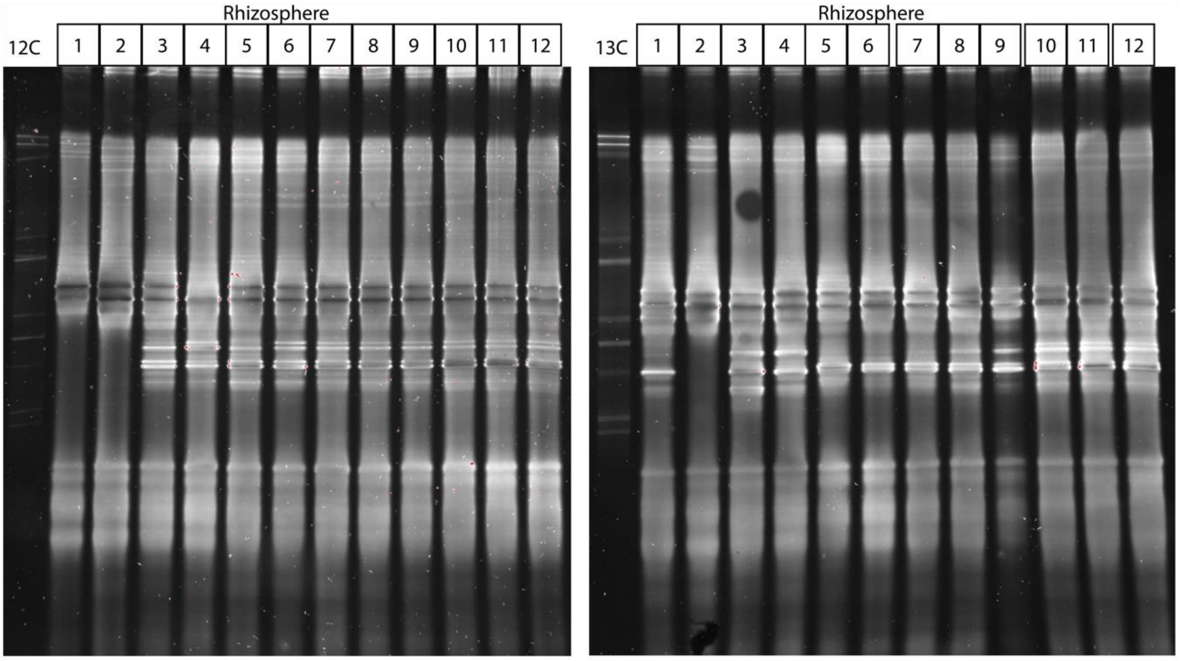


**Supplementary Figure 3**. Denaturing gel gradient electrophoresis (DGGE) showing archaeal 16S rRNA gene diversity across the 12 fractions generated for stable isotope probing for the rhizosphere associated with one $^{12}C$ control (left) and $^{13}C$ labelled (right) plant. Primers indicated in Supplementary Table 6.

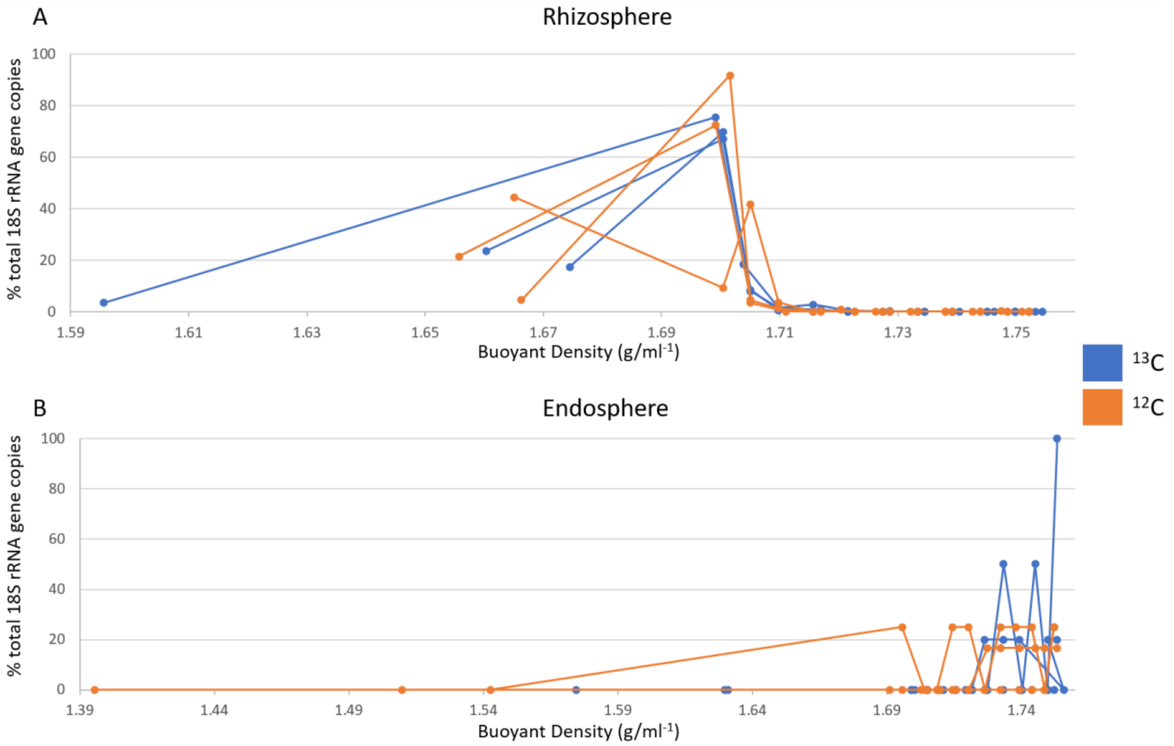


**Supplementary figure 4**. Quantitative PCR against the fungal 18S rRNA gene to test for $^{13}C$ labelling of the fungal community across fractions form the stable isotope probing. Graphs shows the percent of total 18S rRNA genes found within each of the 12 fractions for each plant (plotted as buoyant densities for that fraction in g/ml$^{-1}$) for $^{12}C$ control and $^{13}C$ labelled wheat plants from rhizosphere (A) and endosphere compartments (B) (n=3).

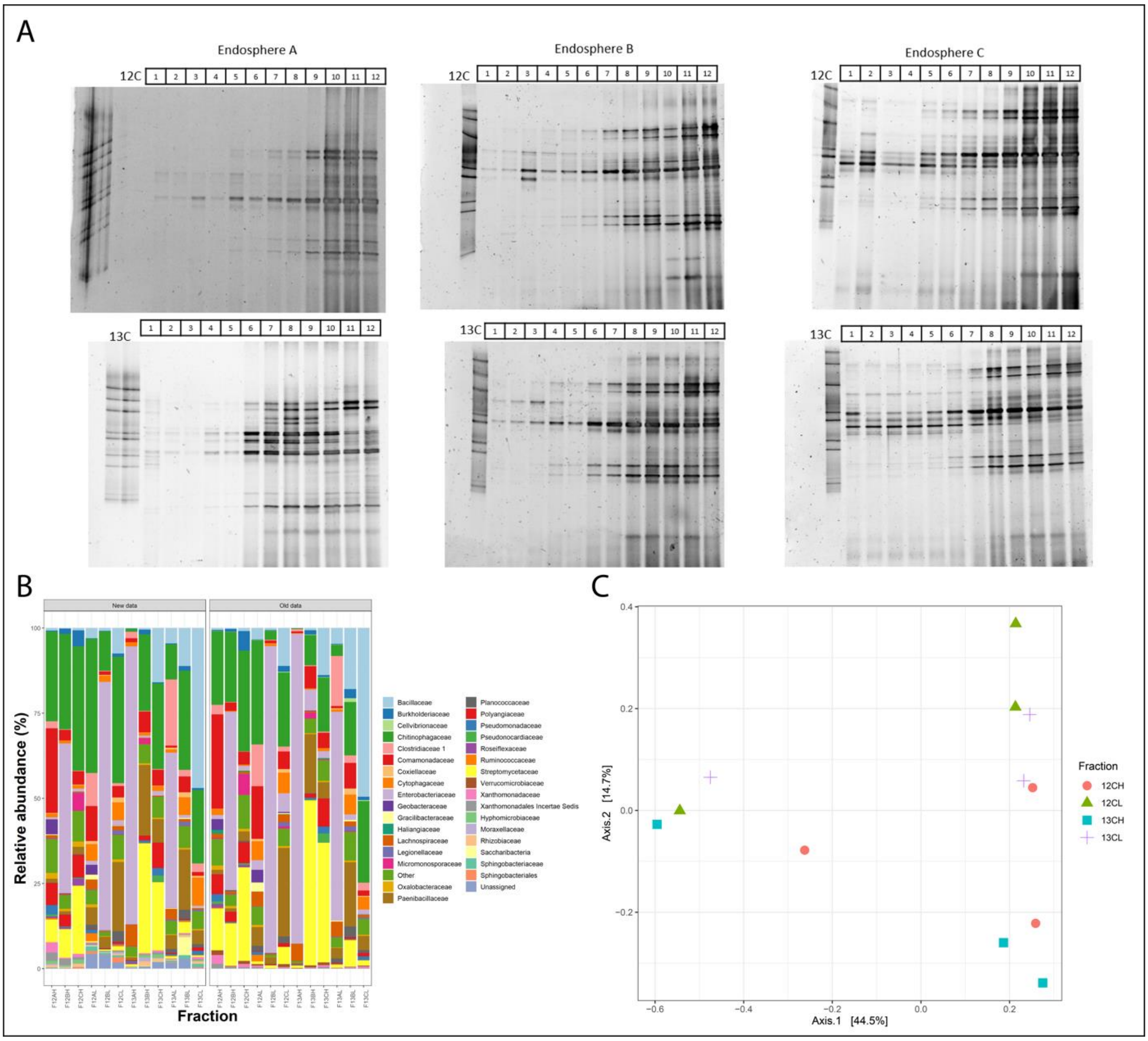


**Supplementary Figure 5.** Endosphere stable isotope probing data. **A** Denaturing gel gradient electrophoresis (DGGE) showing bacterial 16S rRNA gene diversity across the 12 fractions generated for stable isotope probing for the endosphere associated with three $^{12}$C control (top) and $^{13}$C labelled (bottom) plants. These gels show a shift in the bacterial community towards the heavy fraction of $^{13}$C labelled plants. **B** Bars show the relative abundance of each bacterial group within the pooled sequenced $^{12}$C heavy, $^{12}$C light, $^{13}$C heavy and $^{13}$C light fractions (n=3), for two separate sequencing runs on the same samples (old & new). **C** Principle coordinates analysis (PCoA) on bray cutis dissimilarities for the endosphere $^{12}$C heavy (orange/circle), $^{12}$C light (green/triangle), $^{13}$C heavy (blue/square) and $^{13}$C light (purple/cross) fractions (n=3).

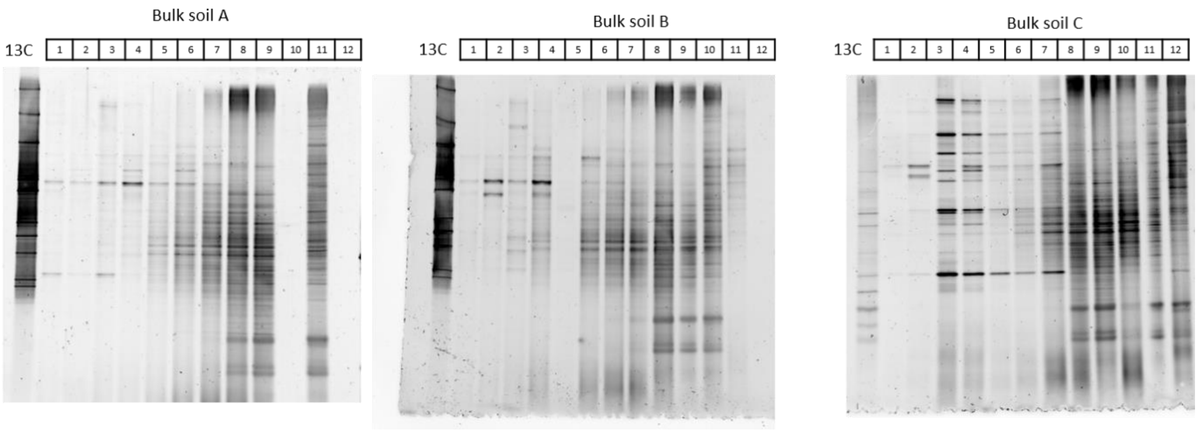


**Supplementary Figure 6**. Denaturing gel gradient electrophoresis (DGGE) showing bacterial 16S rRNA gene diversity across the 12 fractions generated for stable isotope probing for the $^{13}C$ unplanted soil controls (N=3). Primers indicated in Supplementary Table 6.

Supplementary Table 1. SIMPER outputs

| **Pot grown wheat, all soil types** | | | | | |
|---|---|---|---|---|---|
| **Comparison- Bulk soil-Endosphere** | | | **Bulk soil-Rhizosphere** | | |
| **OTU** | **Percent contribution** | **p-value** | **OTU** | **Percent contribution** | **p-value** |
| *Streptomycetaceae* | 14.6 | 0.0099 | *Burkholderiaceae* | 8.9 | 0.4851 |
| *Burkholderiaceae* | 10.1 | 0.0099 | *Rhizobiaceae* | 5.4 | 0.0099 |
| *Acidobacteria* Subgroup 6 | 3.6 | 0.0099 | *Acidobacteria* Subgroup 6 | 4 | 0.0198 |
| *Chitinophagaceae* | 3.5 | 0.0792 | *Bacillaceae* | 3.4 | 0.0594 |
| *Bacillaceae* | 2.8 | 0.0099 | *Micrococcaceae* | 3.1 | 0.0297 |
| *Xanthobacteraceae* | 2.6 | 0.0099 | *Chitinophagaceae* | 2.6 | 0.9108 |
| *Rhizobiaceae* | 2.4 | 0.6733 | *Xanthobacteraceae* | 2.5 | 0.7227 |
| Solirubrobacterales | 2 | 0.0099 | *Saccharimonadale* | 2.4 | 0.3168 |
| Saccharimonadales | 2 | 0.0792 | *Rubritaleaceae* | 2.3 | 0.0099 |
| *Methyloligellaceae* | 1.6 | 0.0099 | *Pseudomonadaceae* | 2.2 | 0.0099 |
| **Pot grown wheat, all soil types** | | | **Field-grown wheat** | | |
| **Comparison- Rhizosphere-Endosphere** | | | **Comparison- Bulk soil-Endosphere** | | |
| **OTU** | **Percent contribution** | **p-value** | **OTU** | **Percent contribution** | **p-value** |
| *Streptomycetaceae* | 16.7 | 0.0099 | *Streptomycetaceae* | 16.39547552 | 0.00990 |
| *Burkholderiaceae* | 7.2 | 0.7327 | *Burkholderiaceae* | 6.073676003 | 0.00990 |
| *Chitinophagaceae* | 3.8 | 0.3168 | *Firmicutes* | 5.602921016 | 0.00990 |
| *Rhizobiaceae* | 3.3 | 0.4752 | *Acidobacteria* Subgroup 6 | 5.547716313 | 0.00990 |
| *Micrococcaceae* | 3 | 0.0099 | *Sphingobacteriaceae* | 4.910748857 | 0.04950 |
| *Sphingobacteriaceae* | 2.4 | 0.1584 | *Bacillaceae* | 4.262550068 | 0.00990 |
| *Xanthobacteraceae* | 2.1 | 0.8416 | *Methyloligellaceae* | 2.413515939 | 0.00990 |

| | | | | | |
|---|---|---|---|---|---|
| Solirubrobacterales | 2.1 | 0.2277 | *Promicromonosporaceae* | 2.307694859 | 0.01980 |
| *Acidobacteria* Subgroup 6 | 2 | 1 | Solirubrobacterales | 2.038018274 | 0.00990 |
| **Field-grown wheat** | | | | | |
| **Bulk soil-Rhizosphere** | | | **Comparison- Rhizosphere-Endosphere** | | |
| **OTU** | **Percent contribution** | **p-value** | **OTU** | **Percent contribution** | **p-value** |
| *Sphingobacteriaceae* | 7.82061 | 0.10891 | *Streptomycetaceae* | 20.89116 | 0.01980 |
| *Burkholderiaceae* | 5.59509 | 0.77227 | *Firmicutes* | 7.499111 | 0.01980 |
| *Micrococcaceae* | 4.89642 | 0.10891 | *Micrococcaceae* | 4.78363 | 0.00990 |
| *Acidobacteria* Subgroup 6 | 4.74396 | 0.93069 | *Acidobacteria* Subgroup 6 | 4.240018 | 0.35643 |
| *Bacillaceae* | 4.08710 | 0.60396 | *Burkholderiaceae* | 4.209923 | 0.65346 |
| *Pseudomonadaceae* | 3.61463 | 0.00990 | *Sphingobacteriaceae* | 3.371828 | 0.98019 |
| *Rhizobiaceae* | 3.12084 | 0.00990 | *Bacillaceae* | 2.840192 | 0.68316 |
| *Methyloligellaceae* | 3.02557 | 0.17821 | *Pseudomonadaceae* | 2.130237 | 0.29703 |
| *Spirosomaceae* | 2.78842 | 0.17821 | *Promicromonosporaceae* | 2.044043 | 0.34653 |
| Solirubrobacterales | 2.42232 | 0.17821 | *Xanthobacteraceae* | 1.735366 | 0.02970 |

Supplementary Table 2. PERMANOVA results

| **Stem elongation compared to pot agricultural** | | | |
|---|---|---|---|
| **Community = Archaea** | | | |
| **Compartment** | **Permutations** | **$R^2$** | **p-value** |
| Bulk Soil | 999 | 0.12 | 0.7 |
| Rhizosphere | 999 | 0.12 | 0.7 |
| Endosphere | 999 | 0.34 | 0.2 |
| **Community = Bacteria** | | | |
| Bulk Soil | 999 | 0.3 | 0.1 |
| Rhizosphere | 999 | 0.58 | 0.1 |
| Endosphere | 999 | 0.53 | 0.1 |
| **Community = Fungi** | | | |
| Bulk Soil | Fungi | 0.53 | 0.1 |
| Rhizosphere | 999 | 0.39 | 0.1 |
| Endosphere | 999 | 0.42 | 0.1 |

| **Test for effect of root compartment on senescent communities** | | | |
|---|---|---|---|
| **Community** | **Permutations** | **$R^2$** | **p-value** |
| Archaea | 999 | 0.68 | 0.01 |
| Bacteria | 999 | 0.74 | 0.005 |
| Fungi | 999 | 0.73 | 0.005 |
| **Test for effect of soil type on community composition** | | | |
| **Bulk Soil** | | | |
| Archaea | 999 | 0.94 | 0.003 |
| Bacteria | 999 | 0.87 | 0.001 |
| Fungi | 999 | 0.81 | 0.004 |
| **Rhizosphere** | | | |
| Archaea | 999 | 0.97 | 0.004 |
| Bacteria | 999 | 0.83 | 0.001 |
| Fungi | 999 | 0.66 | 0.004 |
| **Endosphere** | | | |
| Archaea | 999 | 0.87 | 0.004 |
| Bacteria | 999 | 0.6 | 0.001 |

Supplementary Table 3. Soil chemical properties

| Parameter | Measurement | SAC Rating |
|---|---|---|
| **Agricultural Soil, John Innes Centre Field Studies Site (sampled 18 04 2019)** | | |
| pH | 7.97 | n/a |
| Phosphorus (mg/kg) | 81.63 | High |
| Potassium (mg/kg) | 103 | Moderate |
| Magnesium (mg/kg) | 34.8 | Very Low |
| Nitrate (g/kg) | 73.92 | n/a |
| Ammonium (g/kg) | 6.97 | n/a |
| Organic matter (%) | 2.26 | n/a |
| **F2 Levington Compost** | | |
| pH | 4.98 | n/a |
| Phosphorus (mg/kg) | 880.5 | Excessively high |
| Potassium (mg/kg) | 2508 | Excessively high |
| Magnesium (mg/kg) | 6021 | Excessively high |
| Nitrate (g/kg) | 5809.49 | n/a |
| Ammonium (g/kg) | 192.18 | n/a |
| Organic matter (%) | 91.08 | n/a |

Supplementary Table 4. SIP fractions used for sequencing

| Origin of fractions | Fraction classification | Pooled fractions | fraction density range (g.ml$^{-1}$) |
|---|---|---|---|
| Endosphere A | 12H | 7,8,9 | 1.7086-1.7204 |
| | 12L | 11,12 | 1.3956-1.6910 |
| | 13H | 7,8,9 | 1.7098-1.7227 |
| | 13L | 11,12 | 1.6004-1.6992 |
| Endosphere B | 12H | 7,8,9 | 1.7098-1.7204 |
| | 12L | 11,12 | 1.5097-1.6957 |
| | 13H | 7,8,9 | 1.7098-1.7216 |
| | 13L | 11,12 | 1.5745-1.6992 |
| Endosphere C | 12H | 7,8,9 | 1.7086-1.7204 |
| | 12L | 11,12 | 1.5427-1.6957 |
| | 13H | 7,8,9 | 1.7110-1.7216 |
| | 13L | 11,12 | 1.6310-1.7004 |
| Rhizosphere A | 12H | 7,8 | 1.7169-1.7227 |
| | 12L | 10,11 | 1.6992-1.7051 |
| | 13H | 7,8 | 1.7169-1.7216 |
| | 13L | 10,11 | 1.6992-1.7039 |
| Rhizosphere B | 12H | 7,8 | 1.7157-1.7204 |
| | 12L | 10,11 | 1.7004-1.7051 |
| | 13H | 7,8 | 1.7157-1.7216 |

| | | | |
|---|---|---|---|
| | 13L | 10,11 | 1.7004-1.7051 |
| Rhizosphere C | 12H | 7,8 | 1.7169-1.7227 |
| | 12L | 10,11 | 1.7016-1.7051 |
| | 13H | 7,8 | 1.7157-1.7216 |
| | 13L | 10,11 | 1.7004-1.7051 |
| Unplanted A | 13H | 8,9 | 1.7086-1.7145 |
| | 13L | 11 | 1.6992 |
| Unplanted B | 13H | 8 | 1.7145 |
| | 13L | 10,11 | 1.7004-1.7039 |
| Unplanted C | 13H | 8,9 | 1.7098-1.7157 |
| | 13L | 11,12 | 1.6745-1.7004 |

Supplementary Table 5. DESeq2 outputs to identify root exudate utilisers in the rhizosphere

| **$^{12}$C heavy compared to $^{13}$C heavy** | | | | |
|---|---|---|---|---|
| **OTU** | **baseMean** | **log2FoldChange** | **lfsSE *** | **Padj** |
| *Enterobacteriaceae* | 1525.585247 | 9.225478608 | 1.330868073 | 8.30E-11 |
| *Paenibacillaceae* | 542.7147998 | 6.131222282 | 0.71096459 | 1.62E-16 |
| *Verrucomicrobiaceae* | 1540.476332 | 5.525652719 | 0.38986191 | 1.34E-43 |
| *Pseudomonadaceae* | 747.4943245 | 5.296358431 | 1.327440714 | 0.000347898 |
| *Oxalobacteraceae* | 1121.798139 | 5.093564169 | 0.466396414 | 3.05E-26 |
| *Cellvibrionaceae* | 143.5596665 | 4.563865623 | 0.692665528 | 6.33E-10 |
| *Comamonadaceae* | 1965.095913 | 4.553967836 | 0.399631593 | 2.20E-28 |
| *Fibrobacteraceae* | 308.4266565 | 4.250982079 | 0.691291828 | 9.73E-09 |
| *Rhizobiaceae* | 200.6141532 | 3.586599358 | 0.543771655 | 6.33E-10 |
| *Cytophagaceae* | 333.3918811 | 3.485162531 | 0.695142566 | 4.45E-06 |
| *Micrococcaceae* | 1002.919707 | 3.188857943 | 0.72279587 | 6.41E-05 |
| *Microbacteriaceae* | 182.5423665 | 2.316546635 | 0.708211293 | 0.004465608 |
| *Xanthomonadaceae* | 191.2474009 | 2.210211572 | 0.38114642 | 6.68E-08 |
| *Intrasporangiaceae* | 196.5957779 | 1.679541697 | 0.522170126 | 0.005191223 |
| *Polyangiaceae* | 174.8113772 | 1.441127742 | 0.521692567 | 0.016393349 |
| **$^{13}$C light compared to $^{13}$C heavy** | | | | |
| **OTU** | **baseMean** | **log2FoldChange** | **lfsSE *** | **Padj** |
| *Enterobacteriaceae* | 1193.557819 | 5.663078218 | 1.200852813 | 1.47E-05 |
| *Paenibacillaceae* | 474.7391477 | 3.450277252 | 0.717541055 | 1.06E-05 |
| *Verrucomicrobiaceae* | 1240.232555 | 4.817514423 | 0.470756764 | 4.58E-23 |
| *Pseudomonadaceae* | 640.932411 | 2.82797024 | 1.152315253 | 0.034597109 |
| *Oxalobacteraceae* | 882.3908862 | 6.866216698 | 0.630972123 | 6.88E-26 |
| *Cellvibrionaceae* | 122.6921288 | 3.359934177 | 0.602489726 | 3.00E-07 |

| | | | | |
|---|---|---|---|---|
| *Comamonadaceae* | 1557.064356 | 4.810895677 | 0.375167998 | 1.19E-35 |
| *Fibrobacteraceae* | 247.2869102 | 3.558583564 | 0.645095913 | 3.77E-07 |
| *Rhizobiaceae* | 194.8884915 | 1.676668472 | 0.519657919 | 0.004548681 |
| *Cytophagaceae* | 283.9658689 | 2.520107112 | 0.595381826 | 0.000113113 |
| *Micrococcaceae* | 918.575397 | 1.673919306 | 0.649494375 | 0.025268353 |
| *Microbacteriaceae* | 144.683441 | 2.15700887 | 0.693944173 | 0.006585131 |
| *Xanthomonadaceae* | 167.3075238 | 1.56520526 | 0.53711052 | 0.011040717 |
| *Intrasporangiaceae* | 158.6061631 | 1.50689272 | 0.517691543 | 0.011040717 |
| *Polyangiaceae* | 119.7725455 | 2.424270358 | 0.530116698 | 2.62E-05 |
| *lfsSE= log2 Fold Change Standard Error | | | | |

Supplementary Table 6. Primers

| **Primer** | **Target** | **Uses** | **Sequence** | **Reference** |
|---|---|---|---|---|
| PRK341F-GC | Bacterial 16S | DGGE PCR Amplification | CGCCCGCCGCGCGCGGCGGGC GGGGCGGGGGCACGGGGGGCC TACGGGAGGCAGCAG | (Muyzer *et al.*, 1993) [94] |
| 518R | Bacterial 16S | DGGE PCR Amplification | ATTACCGCGGCTGCTGG | (Muyzer *et al.*, 1993) [94] |
| A771F-GC | Archaeal 16S | DGGE PCR Amplification | CGCCCGCCGCGCGCGGCGGG CGGGGCGGGGGCACGGGGGG ACGGTGAGGGATGAAAGCT | (Ochsenreiter *et al.*, 2003) [95] |
| A957R | Archaeal 16S | DGGE PCR Amplification, qPCR | CGGCGTTGACTCCAATTG | (Ochsenreiter *et al.*, 2003) [95] |
| A109F | Archaeal 16S | PCR amplification, qPCR standard amplification | ACKGCTCAGTAACACGT | (Großkopf *et al.*, 1998) [96] |
| A1000R | Archaeal 16S | PCR amplification, qPCR standard amplification | GGCCATGCACYWCYTCTC | (Gantner *et al.*, 2011) [97] |
| PRK341F | Bacterial 16S | PCR amplification/ Sequencing, qPCR standard amplification | CCTACGGGRBGCASCAG | (Yu *et al.*, 2005) [98] |
| MPRK806R | Bacterial 16S | PCR amplification/ Sequencing, qPCR standard amplification | GGACTACNNGGGTATCTAAT | (Yu *et al.*, 2005) [98] |
| fITS7F | Fungal ITS2 | PCR Amplification/ sequencing | GTGARTCATCGAATCTTTG | (Ihrmark *et al.*, 2012) [99] |
| ITS4R_2 | Fungal ITS2 | PCR Amplification/ sequencing | TCCTCCGCTTATTGATATGC | (White *et al.*, 1990) [100] |

| | | | | |
|---|---|---|---|---|
| A771F | Archaeal 16S | qPCR | ACGGTGAGGGATGAAAGCT | (Ochsenreiter *et al.*, 2003) [95] |
| FR1Fw | Fungal 18S | qPCR | AICCATTCAATCGGTAIT | (Vainio *et al.*, 2000) [101] |
| FF390Rev | Fungal 18S | qPCR | CGATAACGAACGAGACCT | (Vainio *et al.*, 2000) [101] |
| Com1F | Bacterial 16S | qPCR | CAGCAGCCGCGGTAATAC | (Fredriksson *et al.*, 2013) [102] |
| 769R | Bacterial 16S | qPCR | ATCCTGTTTGMTMCCCVCRC | (Rastogi *et al.*, 2010) [103] |
| F18SS03-F | Fungal 18S | qPCR standard amplification | AGATCCTGAGGCCTCACTA | This Study |
| F18SS03-R | Fungal 18S | qPCR standard amplification | GCCGTTCTTAGTTGGTGGAG | This Study |
| A0349F | Archaeal 16S | Sequencing | GYGCASCAGKCGMGAAW | (Takai *et al.*, 2000) [104] |
| A0519R | Archaeal 16S | Sequencing | TTACCGCGGCKGCTG | (Takai *et al.*, 2000) [104] |
| CrenamoA23f | Archaeal *amoA* | DGGE | ATGGTCTGGCTWAGACG | (Tourna *et al.*, 2008) [105] |
| CrenamoA616r | Archaeal *amoA* | DGGE | GCCATCCATCTGTATGTCCA | (Tourna *et al.*, 2008) [105] |

Supplementary Table 7. PCR Conditions for metabarcoding, qPCR & DGGE PCRs

| **PCR Component** | **Volume (µl)** |
|---|---|
| 2x PCRBio BioMix™ red, containing BIOTAQ™ DNA Polymerase or 2x PCRBio Ultra mix, containing Ultra DNA Polymerase | 10 |
| Forward or reverse primer (10mM stock) | 1 |
| Template DNA | 2 (DNA extract)<br>1 (Round 1 product for nested PCR) |
| Sterile $dH_2O$ | Up to 20µl |
| **qPCR mix** | |
| 2x SYBR Green Luna® Universal qPCR Master Mix | 10µl |
| Forward or reverse primer (10mM stock) | 0.5µl |
| DNA template (20ng/µl stock) | 5µl |
| Sterile MilliQ $dH_2O$ | Up to 20µl |
| **Thermocycler programs** | |
| A0109F/A1000R<br>Archaeal 16S rRNA gene<br>For sequencing & round one of DGGE | 1. 95°C for 1 minute<br>2. 35x cycles of 95°C for 30 seconds, 59°C for 30 seconds, 72°C for 45 seconds<br>3. 72°C for 1 minute |
| A771F-GC/A957R | 1. 95°C for 1 minute |

| | |
|---|---|
| Archaeal 16S rRNA gene<br>For round two of DGGE | 2. 35 cycles of 94°C for 30 seconds, 55°C for 30 seconds, 72°C for 1 minute<br>3. 72°C for 10 minutes |
| PRK341F/MPRK806R or fITS7F/ITS4R<br>Bacterial 16S rRNA gene or Fungal ITS2 region<br>For sequencing | 1. 95°C for 1 minute<br>2. 30 cycles of 95°C for 15 seconds, 55°C for 15 seconds, 72°C for 15 seconds<br>3. 72°C for 10 minutes |
| A771F/A957R, FR1Fw/FF390Rev or Com1F/769R qPCR assays | 1. 95°C for 10 minutes<br>2. 35 cycles of 95°C for 15 seconds and 60°C annealing/extension/read step for 30 seconds |

Supplementary Table 8. Qiime 2 Dada2 settings

| **Amplicon** | **p-trim-left-f** | **p-trim-left-r** | **p-trunc-len-f/r** |
|---|---|---|---|
| A0349F/A0519R Archaeal 16S | 17 | 15 | 120 |
| PRK341F/ MPRK806R Bacterial 16S | 17 | 20 | 230 |
| fITS7F/ ITS4R Fungal ITS2 | 19 | 20 | 195 |

Supplementary Table 9. Qiime2 taxonomy-based filtering stats

| **Experiment** | **Amplicon** | **Sample** | **Total No. trimmed quality filtered reads** | **No. reads removed by taxonomic filtering** | **No. reads remaining** | **% reads discarded** |
|---|---|---|---|---|---|---|
| Metabarcoding | A0349F/A0519R Archaeal 16S | BS51.A | 68773 | 475 | 68298 | 0.690678 |
| | | BS52.A | 63866 | 625 | 63241 | 0.978611 |
| | | BS53.A | 74447 | 575 | 73872 | 0.772362 |
| | | BSA1.A | 55327 | 867 | 54460 | 1.567047 |
| | | BSA2.A | 54695 | 195 | 54500 | 0.356523 |
| | | BSA3.A | 49970 | 390 | 49580 | 0.780468 |
| | | BSF1.A | 78012 | 1695 | 76317 | 2.172743 |
| | | BSF2.A | 75560 | 1892 | 73668 | 2.50397 |
| | | BSF3.A | 78741 | 1574 | 77167 | 1.998959 |
| | | BSL1.A | 207703 | 11458 | 196245 | 5.516531 |
| | | BSL2.A | 196347 | 8785 | 187562 | 4.474222 |
| | | BSL3.A | 133107 | 4169 | 128938 | 3.132067 |

| | | | | | | |
|---|---|---|---|---|---|---|
| | | E51.A | 93810 | 119 | 93691 | 0.126852 |
| | | E52.A | 92550 | 711 | 91839 | 0.768233 |
| | | E53.A | 109472 | 330 | 109142 | 0.301447 |
| | | EA1.A | 79035 | 54 | 78981 | 0.068324 |
| | | EA2.A | 72672 | 6124 | 66548 | 8.426904 |
| | | EA3.A | 58574 | 34 | 58540 | 0.058046 |
| | | EF1.A | 78178 | 2079 | 76099 | 2.659316 |
| | | EF2.A | 59310 | 442 | 58868 | 0.745237 |
| | | EF3.A | 57468 | 856 | 56612 | 1.489525 |
| | | EL1.A | 126246 | 159 | 126087 | 0.125945 |
| | | EL2.A | 133715 | 200 | 133515 | 0.149572 |
| | | RZ51.A | 69659 | 2320 | 67339 | 3.33051 |
| | | RZ52.A | 75587 | 1549 | 74038 | 2.049294 |
| | | RZ53.A | 68546 | 1174 | 67372 | 1.712718 |
| | | RZA1.A | 58037 | 1202 | 56835 | 2.071093 |
| | | RZA2.A | 90071 | 1176 | 88895 | 1.305637 |
| | | RZA3.A | 68877 | 1513 | 67364 | 2.196669 |
| | | RZF1.A | 37285 | 304 | 36981 | 0.815341 |
| | | RZF2.A | 48495 | 321 | 48174 | 0.661924 |
| | | RZF3.A | 56662 | 657 | 56005 | 1.159507 |
| | | RZL1.A | 125065 | 3873 | 121192 | 3.09679 |
| | | RZL2.A | 125751 | 3886 | 121865 | 3.090234 |
| | | RZL3.A | 123964 | 2723 | 121241 | 2.196605 |
| | | EFS1.A | 239780 | 2790 | 236990 | 1.1 |
| | | EFS2.A | 247035 | 643 | 246392 | 0.260287 |
| | | EFS3.A | 199509 | 1400 | 198109 | 0.701723 |
| | | RZFS1.A | 247152 | 881 | 246271 | 0.356461 |
| | | RZFS2.A | 239414 | 935 | 238479 | 0.390537 |
| | | RZFS3.A | 241137 | 280 | 240857 | 0.116117 |
| | | BSFS1.A | 254643 | 226 | 254417 | 0.088752 |
| | | BSFS2.A | 304279 | 289 | 303990 | 0.949786 |
| | | BSFS3.A | 478179 | 808 | 477371 | 0.168974 |
| | PRK341F/ MPRK806R Bacterial 16S | B.BS51 | 22970 | 320 | 22650 | 1.393121 |
| | | B.BS52 | 12809 | 510 | 12299 | 3.981575 |
| | | B.BS53 | 22638 | 396 | 22242 | 1.749271 |
| | | B.BSA1 | 13386 | 455 | 12931 | 3.399074 |
| | | B.BSA2 | 18260 | 778 | 17482 | 4.260679 |
| | | B.BSA3 | 17575 | 977 | 16598 | 5.559033 |
| | | B.BSL1 | 32363 | 3322 | 29041 | 10.26481 |
| | | B.BSL2 | 28475 | 3580 | 24895 | 12.57243 |

| | | | | | | |
|---|---|---|---|---|---|---|
| | | B.BSL3 | 28042 | 3478 | 24564 | 12.40282 |
| | | B.E51 | 51847 | 50333 | 1514 | 97.07987 |
| | | B.E52 | 46632 | 46145 | 487 | 98.95565 |
| | | B.E53 | 49496 | 46634 | 2862 | 94.21771 |
| | | B.EA1 | 46470 | 42828 | 3642 | 92.16269 |
| | | B.EA2 | 49475 | 47264 | 2211 | 95.53108 |
| | | B.EA3 | 39328 | 32351 | 6977 | 82.25946 |
| | | B.EL1 | 40368 | 39732 | 636 | 98.42449 |
| | | B.EL2 | 46651 | 46125 | 526 | 98.87248 |
| | | B.EL3 | 48154 | 46843 | 1311 | 97.27748 |
| | | B.RZ51 | 17754 | 976 | 16778 | 5.497353 |
| | | B.RZ52 | 18840 | 983 | 17857 | 5.217622 |
| | | B.RZ53 | 17688 | 782 | 16906 | 4.421076 |
| | | B.RZA1 | 22737 | 687 | 22050 | 3.021507 |
| | | B.RZA2 | 23577 | 813 | 22764 | 3.448276 |
| | | B.RZA3 | 20817 | 647 | 20170 | 3.108037 |
| | | B.RZL1 | 21777 | 632 | 21145 | 2.902144 |
| | | B.RZL2 | 26411 | 1015 | 25396 | 3.843096 |
| | | B.RZL3 | 27594 | 1896 | 25698 | 6.871059 |
| | | B.BSF1 | 17020 | 1101 | 15919 | 6.46886 |
| | | B.BSF2 | 15340 | 568 | 14772 | 3.702738 |
| | | B.BSF3 | 19415 | 414 | 19001 | 2.132372 |
| | | B.EF1 | 41256 | 24671 | 16585 | 59.79979 |
| | | B.EF2 | 42388 | 27888 | 14500 | 65.79221 |
| | | B.EF3 | 44801 | 27308 | 17493 | 60.954 |
| | | B.RZF1 | 21359 | 1201 | 20158 | 5.622922 |
| | | B.RZF2 | 24495 | 472 | 24023 | 1.926924 |
| | | B.RZF3 | 19582 | 760 | 18822 | 3.881115 |
| | | B.EFS1 | 53449 | 138 | 53311 | 0.25819 |
| | | B.EFS2 | 60192 | 350 | 59842 | 0.581473 |
| | | B.EFS3 | 54148 | 755 | 53393 | 1.394327 |
| | | B.RZFS1 | 47685 | 367 | 47318 | 0.769341 |
| | | B.RZFS2 | 46770 | 2122 | 44648 | 4.537096 |
| | | B.RZFS3 | 48877 | 844 | 48033 | 1.726784 |
| | | B.BSFS1 | 43114 | 7244 | 35870 | 16.80197 |
| | | B.BSFS2 | 42848 | 4569 | 38279 | 10.66327 |
| | | B.BSFS3 | 48794 | 6410 | 42384 | 13.13686 |
| | fITS7F/ ITS4R Fungal ITS2 | F.BS51 | 41729 | n/a | n/a | n/a |
| | | F.BS52 | 45565 | n/a | n/a | n/a |

| | | | | | | |
|---|---|---|---|---|---|---|
| | | F.BS53 | 46115 | n/a | n/a | n/a |
| | | F.BSA1 | 75779 | n/a | n/a | n/a |
| | | F.BSA2 | 100787 | n/a | n/a | n/a |
| | | F.BSA3 | 94052 | n/a | n/a | n/a |
| | | F.BSF1 | 129866 | n/a | n/a | n/a |
| | | F.BSF2 | 100424 | n/a | n/a | n/a |
| | | F.BSF3 | 81109 | n/a | n/a | n/a |
| | | F.BSL1 | 98179 | n/a | n/a | n/a |
| | | F.BSL2 | 131559 | n/a | n/a | n/a |
| | | F.BSL3 | 120348 | n/a | n/a | n/a |
| | | F.E51 | 101772 | n/a | n/a | n/a |
| | | F.E52 | 114929 | n/a | n/a | n/a |
| | | F.E53 | 44992 | n/a | n/a | n/a |
| | | F.EA1 | 75592 | n/a | n/a | n/a |
| | | F.EA2 | 65668 | n/a | n/a | n/a |
| | | F.EA3 | 89945 | n/a | n/a | n/a |
| | | F.EF1 | 89057 | n/a | n/a | n/a |
| | | F.EF2 | 69637 | n/a | n/a | n/a |
| | | F.EF3 | 102551 | n/a | n/a | n/a |
| | | F.RZ51 | 84654 | n/a | n/a | n/a |
| | | F.RZ52 | 37011 | n/a | n/a | n/a |
| | | F.RZ53 | 31117 | n/a | n/a | n/a |
| | | F.RZA1 | 102095 | n/a | n/a | n/a |
| | | F.RZA2 | 71665 | n/a | n/a | n/a |
| | | F.RZA3 | 84015 | n/a | n/a | n/a |
| | | F.RZF1 | 51408 | n/a | n/a | n/a |
| | | F.RZF2 | 99541 | n/a | n/a | n/a |
| | | F.RZF3 | 45291 | n/a | n/a | n/a |
| | | F.RZL1 | 110595 | n/a | n/a | n/a |
| | | F.RZL2 | 97230 | n/a | n/a | n/a |
| | | F.RZL3 | 81901 | n/a | n/a | n/a |
| | | F.EFS1 | 129650 | n/a | n/a | n/a |
| | | F.EFS2 | 70055 | n/a | n/a | n/a |
| | | F.EFS3 | 76512 | n/a | n/a | n/a |
| | | F.RZFS1 | 56010 | n/a | n/a | n/a |
| | | F.RZFS2 | 64946 | n/a | n/a | n/a |
| | | F.RZFS3 | 36457 | n/a | n/a | n/a |
| | | F.BSFS1 | 104076 | n/a | n/a | n/a |
| | | F.BSFS2 | 71894 | n/a | n/a | n/a |

| | | | | | | |
|---|---|---|---|---|---|---|
| | | F.BSFS3 | 80601 | n/a | n/a | n/a |
| Endosphere SIP sequencing (1st run) | PRK341F/ MPRK806R Bacterial 16S | 12CHa | 93805 | 43076 | 50729 | 45.92079 |
| | | 12CLa | 116684 | 89134 | 27550 | 76.38922 |
| | | 13CHa | 108182 | 29914 | 78268 | 27.65155 |
| | | 13CLa | 97947 | 48972 | 48975 | 49.99847 |
| | | 12CHb | 108089 | 43142 | 64947 | 39.9134 |
| | | 12CLb | 105021 | 50010 | 55011 | 47.61905 |
| | | 13CHb | 95683 | 73700 | 21983 | 77.02518 |
| | | 13CLb | 71120 | 52215 | 18905 | 73.41817 |
| | | 12CHb | 100291 | 56735 | 43556 | 56.57038 |
| | | 12CLb | 92092 | 84369 | 7723 | 91.61382 |
| | | 13CHb | 83840 | 46040 | 37800 | 54.91412 |
| | | 13CLb | 85336 | 45571 | 39765 | 53.40185 |
| Rhizosphere and Bulk soil SIP sequencing | | 12CHa | 13421 | 885 | 12536 | 6.594144 |
| | | 12CLa | 12970 | 798 | 12172 | 6.15266 |
| | | 13CHa | 15960 | 336 | 15624 | 2.105263 |
| | | 13CLa | 12166 | 142 | 12024 | 1.167187 |
| | | 12CHb | 22378 | 3195 | 19183 | 14.27742 |
| | | 12CLb | 14140 | 986 | 13154 | 6.973126 |
| | | 13CHb | 14082 | 321 | 13761 | 2.279506 |
| | | 13CLb | 10732 | 286 | 10446 | 2.664927 |
| | | 12CHc | 15785 | 316 | 15469 | 2.001901 |
| | | 12CLc | 8832 | 380 | 8452 | 4.302536 |
| | | 13CHc | 12383 | 172 | 12211 | 1.389001 |
| | | 13CLc | 10651 | 146 | 10505 | 1.370763 |
| | | 13CHa | 19395 | 4230 | 15165 | 21.80974 |
| | | 13CLa | 14826 | 1932 | 12894 | 13.03116 |
| | | 13CHb | 19529 | 1409 | 18120 | 7.214911 |
| | | 13CLb | 17501 | 1740 | 15761 | 9.942289 |
| | | 13CHc | 15543 | 2230 | 13313 | 14.34729 |
| | | 13CLc | 18138 | 2585 | 15553 | 14.25185 |
| Endosphere SIP sequencing (2nd run) | | 12CHa | 129499 | 76038 | 53461 | 41.28294 |
| | | 12CLa | 112494 | 35149 | 77345 | 68.75478 |
| | | 12CHb | 129363 | 68584 | 60779 | 46.9833 |
| | | 12CLb | 128990 | 61089 | 67901 | 52.64051 |
| | | 12CHc | 111366 | 56655 | 54711 | 49.1272 |
| | | 12CLc | 106026 | 27714 | 78312 | 73.86113 |
| | | 13CHa | 145802 | 94298 | 51504 | 35.32462 |
| | | 13CLa | 102901 | 53396 | 49505 | 48.10935 |
| | | 13CHb | 104884 | 40712 | 64172 | 61.18378 |

| | | | | | | |
|---|---|---|---|---|---|---|
| | | 13CLb | 92606 | 31200 | 61406 | 66.30888 |
| | | 13CHc | 124416 | 57972 | 66444 | 53.40471 |
| | | 13CLc | 109262 | 53343 | 55919 | 51.17882 |
| | | **Key**: BS = Bulk Soil, RZ = Rhizosphere, E = Endosphere 5 = 50:50 Mix, A = Agricltural Pot, F = Stem Elongation Field, L = Levington F2 Compost, FS = Senescent Field 1/2/3 = sample replicate, A./B./F./ = Archaeal/Bacterial/Fungal amplicon, 12CL – 12C light, 12CH – 12C heavy, 13CL – 13C light, 13CH – 13C heavy, a/b/c – sample replicate. | | | | |

Supplementary Table 10. DESeq2 output identifying $CO_2$-fixing autotrophs from unplanted soil

| **OTU** | **baseMean** | **log2FoldChange** | **lfsSE *** | **padj** |
|---|---|---|---|---|
| *Gaiellaceae* | 523.2173795 | 2.423212929 | 0.588347809 | 0.000181456 |
| *Gemmatimonadaceae* | 463.0262448 | 2.071076321 | 0.535193797 | 0.000453911 |
| *Acidimicrobiaceae* | 366.6391772 | 1.760321921 | 0.508212808 | 0.001886839 |
| *Micromonosporaceae* | 152.0272389 | 1.923264409 | 0.628460704 | 0.005528292 |
| *Solirubrobacteraceae* | 114.7633207 | 1.91416848 | 0.571306195 | 0.002444309 |
| *Intrasporangiaceae* | 111.1246338 | 2.435159256 | 0.928676387 | 0.016801645 |
| *lfsSE= log2 Fold Change Standard Error | | | | |

Supplementary Table 11. DESeq2 outputs stem elongation field grown wheat- Significantly differentially abundant bacterial taxa

| **Comparison - Bulk Soil-Rhizosphere** | | | | |
|---|---|---|---|---|
| **OTU** | **Base Mean** | **log2 Fold Change** | **lfcSE *** | **padj** |
| *Bacillaceae* | 372.2125 | -0.14622 | 0.239322 | 0.693852 |
| *Bacteriovoracaceae* | 13.98168 | 2.230435 | 0.619679 | 0.002832 |
| *Burkholderiaceae* | 1021.925 | 0.813465 | 0.288343 | 0.029905 |
| *Caulobacteraceae* | 110.498 | 1.628402 | 0.494308 | 0.00759 |
| *Chitinophagaceae* | 165.0146 | 0.94339 | 0.293433 | 0.009317 |
| *Devosiaceae* | 101.8837 | 2.40246 | 0.519061 | 0.000123 |
| *Gaiellaceae* | 258.2943 | -0.38662 | 0.113536 | 0.006009 |
| *Geminicoccaceae* | 26.37856 | -0.35258 | 0.514868 | 0.640873 |
| *Haliangiaceae* | 26.72952 | -0.06375 | 0.511428 | 0.948208 |
| *Hymenobacteraceae* | 24.41687 | -0.34667 | 0.477006 | 0.631582 |
| *Ilumatobacteraceae* | 104.5866 | -0.42953 | 0.354796 | 0.383108 |
| *Microbacteriaceae* | 280.9652 | 0.790671 | 0.218822 | 0.003023 |
| *Micromonosporaceae* | 70.07192 | -0.09121 | 0.283765 | 0.849883 |

| | | | | |
|---|---|---|---|---|
| *Mycobacteriaceae* | 72.29347 | 0.013391 | 0.291328 | 0.982999 |
| *Nitrosomonadaceae* | 34.0193 | -0.54395 | 0.5127 | 0.435947 |
| *Nitrospiraceae* | 34.26428 | -1.08858 | 0.331452 | 0.006069 |
| *Opitutaceae* | 11.38435 | 1.069774 | 0.522292 | 0.094273 |
| *Paenibacillaceae* | 373.3526 | 0.511014 | 0.16033 | 0.009575 |
| *Pedosphaeraceae* | 24.25692 | -1.34006 | 0.45871 | 0.016595 |
| *Planococcaceae* | 52.96895 | -0.19828 | 0.286047 | 0.640873 |
| *Promicromonosporaceae* | 141.5928 | 1.937938 | 0.499305 | 0.001299 |
| *Pseudomonadaceae* | 400.3107 | 1.768233 | 0.417393 | 0.000454 |
| *Pseudonocardiaceae* | 34.84908 | -0.24604 | 0.541155 | 0.773224 |
| *Rhizobiaceae* | 393.09 | 1.211811 | 0.182278 | 2.97E-09 |
| *Roseiflexaceae* | 108.7535 | 0.144605 | 0.398015 | 0.823412 |
| *Rubritaleaceae* | 100.1947 | -0.89572 | 0.244092 | 0.002699 |
| Saccharimonadaceae | 26.26498 | 1.690537 | 0.510286 | 0.00759 |
| Solirubrobacteraceae | 129.6696 | -0.37406 | 0.141224 | 0.044888 |
| Sphingobacteriaceae | 776.3987 | 1.982141 | 0.452171 | 0.000292 |
| Spirosomaceae | 257.6693 | 2.757245 | 0.49784 | 1.53E-06 |
| Streptomycetaceae | 380.0181 | 0.757702 | 0.323487 | 0.076663 |
| Xanthobacteraceae | 252.9046 | -0.04345 | 0.295926 | 0.939636 |
| **Comparison – Rhizosphere-Endosphere** | | | | |
| **OTU** | **Base Mean** | **log2 Fold Change** | **lfcSE*** | **padj** |
| Bacillaceae | 372.2125 | -0.14622 | 0.239322 | 0.693852 |
| Bacteriovoracaceae | 13.98168 | 2.230435 | 0.619679 | 0.002832 |
| Burkholderiaceae | 2152.24 | 1.142141 | 0.254425 | 0.000102 |
| Caulobacteraceae | 326.3971 | 1.281452 | 0.530015 | 0.053849 |
| Chitinophagaceae | 306.5415 | 0.988075 | 0.277398 | 0.002832 |
| Devosiaceae | 310.9739 | 1.247119 | 0.265459 | 4.38E-05 |
| Gaiellaceae | 65.93289 | -0.57671 | 0.483931 | 0.388951 |
| Geminicoccaceae | 26.37856 | -0.35258 | 0.514868 | 0.640873 |
| Haliangiaceae | 26.72952 | -0.06375 | 0.511428 | 0.948208 |
| Hymenobacteraceae | 24.41687 | -0.34667 | 0.477006 | 0.631582 |
| Ilumatobacteraceae | 104.5866 | -0.42953 | 0.354796 | 0.383108 |
| Microbacteriaceae | 364.1172 | 0.699782 | 0.316859 | 0.068361 |
| Micromonosporaceae | 243.0461 | 1.932803 | 0.296194 | 3.39E-09 |
| Mycobacteriaceae | 72.29347 | 0.013391 | 0.291328 | 0.982999 |
| Nitrosomonadaceae | 34.0193 | -0.54395 | 0.5127 | 0.435947 |
| Nitrospiraceae | 34.26428 | -1.08858 | 0.331452 | 0.006069 |
| Opitutaceae | 11.38435 | 1.069774 | 0.522292 | 0.094273 |
| Paenibacillaceae | 367.5622 | 0.532595 | 0.241368 | 0.068361 |
| Pedosphaeraceae | 24.25692 | -1.34006 | 0.45871 | 0.016595 |
| Planococcaceae | 52.96895 | -0.19828 | 0.286047 | 0.640873 |
| Promicromonosporaceae | 631.1794 | 1.583836 | 0.370707 | 0.000242 |

| | | | | |
|---|---|---|---|---|
| Pseudomonadaceae | 337.4026 | -0.05043 | 0.471101 | 0.952873 |
| Pseudonocardiaceae | 314.0513 | 2.733577 | 0.434434 | 7.82E-09 |
| Rhizobiaceae | 503.9776 | 0.568904 | 0.274098 | 0.090323 |
| Roseiflexaceae | 108.7535 | 0.144605 | 0.398015 | 0.823412 |
| Rubritaleaceae | 140.9361 | 0.749244 | 0.331561 | 0.062729 |
| Saccharimonadaceae | 115.8647 | 1.595737 | 0.55365 | 0.017169 |
| Solirubrobacteraceae | 37.03626 | -0.3467 | 0.346116 | 0.465436 |
| Sphingobacteriaceae | 1388.722 | 0.797741 | 0.338137 | 0.054036 |
| Spirosomaceae | 607.5499 | 1.017229 | 0.414429 | 0.052248 |
| Streptomycetaceae | 4613.887 | 2.536194 | 0.400799 | 7.82E-09 |
| Xanthobacteraceae | 252.9046 | -0.04345 | 0.295926 | 0.939636 |
| *lfsSE= log2 Fold Change Standard Error | | | | |

Supplementary Table 12. DESeq2 outputs all samples (compost, agricultural pot, 50:50 mix and field stem elongation)- Significantly differentially bacterial abundant taxa

| **Comparison - Bulk Soil-Rhizosphere** | | | | |
|---|---|---|---|---|
| **OTU** | **Base Mean** | **log2 Fold Change** | **lfcSE *** | **padj** |
| Unknowns | 2068.626 | -1.19715 | 0.411885 | 0.026106 |
| *Burkholderiaceae* | 985.327 | 3.179403 | 0.485224 | 2.83E-09 |
| Uncultured | 756.9371 | -1.25519 | 0.465714 | 0.046897 |
| Unassigned | 638.8419 | -1.13566 | 0.296593 | 0.001838 |
| *Rhizobiaceae* | 308.3923 | 1.995953 | 0.319026 | 9.85E-09 |
| *Streptomycetaceae* | 292.7344 | 1.160709 | 0.344891 | 0.008221 |
| *Pseudomonadaceae* | 267.3359 | 4.65929 | 0.654589 | 1.10E-10 |
| *Rubritaleaceae* | 184.9994 | 3.54871 | 0.808995 | 0.00023 |
| env.OPS 17 | 136.8828 | -2.13424 | 0.673093 | 0.012669 |
| *Haliangiaceae* | 80.43715 | -0.94404 | 0.278696 | 0.008221 |
| *Spirosomaceae* | 59.57213 | 4.777798 | 0.742664 | 4.16E-09 |
| *Pyrinomonadaceae* | 58.62283 | -2.1681 | 0.648117 | 0.008221 |
| *Fibrobacteraceae* | 57.28673 | 2.684165 | 0.854304 | 0.012911 |
| metagenome | 54.20017 | -1.47081 | 0.382967 | 0.001838 |
| *Cellvibrionaceae* | 25.5762 | 2.138713 | 0.668686 | 0.012564 |
| **Comparison – Rhizosphere-Endosphere** | | | | |
| **OTU** | **Base Mean** | **log2 Fold Change** | **lfcSE*** | **padj** |
| *Streptomycetaceae* | 1540.243 | -4.577 | 0.532667 | 7.14E-16 |
| *Erysipelotrichaceae* | 7.280075 | -6.11435 | 0.737884 | 4.91E-15 |
| *Thermomonosporaceae* | 14.85748 | -4.76837 | 0.696907 | 2.18E-10 |
| *Pseudonocardiaceae* | 88.20983 | -4.75912 | 0.809506 | 8.67E-08 |
| *Pedosphaeraceae* | 14.18555 | 2.540781 | 0.498724 | 5.87E-06 |
| *Polyangiaceae* | 161.9785 | -3.4803 | 0.769874 | 8.63E-05 |
| *Chitinophagaceae* | 424.247 | -3.02177 | 0.705147 | 0.000192 |

| | | | | |
|---|---|---|---|---|
| *Rhizobiaceae* | 267.3061 | -1.11239 | 0.258018 | 0.000192 |
| *Promicromonosporaceae* | 88.51291 | -3.44988 | 0.844883 | 0.000414 |
| Unassigned | 78.08997 | 1.719875 | 0.424449 | 0.000427 |
| *Gemmatimonadaceae* | 70.10253 | 2.198493 | 0.561945 | 0.000698 |
| *Solirubrobacteraceae* | 19.04678 | 1.762621 | 0.490947 | 0.002313 |
| *Paenibacillaceae* | 123.1825 | -2.2608 | 0.650672 | 0.003306 |
| WD2101 soil group | 30.91117 | 2.731383 | 0.849034 | 0.007771 |
| *Cellvibrionaceae* | 27.81366 | -2.00047 | 0.628252 | 0.008129 |
| *Cytophagaceae* | 14.87105 | -1.99234 | 0.644695 | 0.010495 |
| *Microscillaceae* | 152.3535 | -2.09948 | 0.719648 | 0.017442 |
| uncultured actinobacterium | 8.533708 | 1.928468 | 0.684966 | 0.022733 |
| *Saccharimonadaceae* | 21.08344 | -2.26986 | 0.819863 | 0.02489 |
| CPla-3 termite group | 11.36798 | 2.312846 | 0.905758 | 0.044792 |
| *lfsSE= log2 Fold Change Standard Error | | | | |

Supplementary Table 13. DESeq2 outputs senescent plants- Significantly differentially abundant taxa

| **Comparison - Bulk Soil-Rhizosphere** | | | | |
|---|---|---|---|---|
| **OTU** | **Base Mean** | **log2 Fold Change** | **lfcSE*** | **padj** |
| *Bacillaceae* | 767.3759 | 0.075595 | 0.050122 | 0.26299 |
| *Bacteriovoracaceae* | 355.3372 | 0.446428 | 0.06923 | 1.13E-08 |
| *Burkholderiaceae* | 1720.065 | 0.178619 | 0.035847 | 1.25E-05 |
| *Caulobacteraceae* | 161.8836 | -0.19486 | 0.11932 | 0.227669 |
| *Chitinophagaceae* | 2062.64 | 0.028955 | 0.034216 | 0.521632 |
| *Devosiaceae* | 91.78086 | 0.204319 | 0.204316 | 0.466624 |
| *Gaiellaceae* | 390.196 | 0.238407 | 0.092834 | 0.032987 |
| *Geminicoccaceae* | 283.8183 | 0.288585 | 0.099276 | 0.01521 |
| *Gemmataceae* | 304.0407 | 0.205954 | 0.083663 | 0.043212 |
| *Haliangiaceae* | 430.3705 | 0.249691 | 0.071967 | 0.003725 |
| *Ilumatobacteraceae* | 576.283 | 0.137523 | 0.072957 | 0.156406 |
| *Microbacteriaceae* | 235.302 | 0.068233 | 0.097531 | 0.576395 |
| *Micromonosporaceae* | 237.3228 | 0.059733 | 0.090561 | 0.592464 |
| *Mycobacteriaceae* | 211.5452 | -0.04304 | 0.122052 | 0.775018 |
| *Nitrosomonadaceae* | 631.786 | 0.202049 | 0.060474 | 0.005216 |
| *Nitrospiraceae* | 348.0111 | 0.019542 | 0.069559 | 0.812289 |
| *Opitutaceae* | 516.4374 | -0.19252 | 0.062091 | 0.010162 |
| *Paenibacillaceae* | 486.8574 | -0.28646 | 0.072825 | 0.000837 |
| *Pedosphaeraceae* | 445.4718 | 0.147492 | 0.083526 | 0.184347 |
| *Planococcaceae* | 347.0514 | -0.26987 | 0.101567 | 0.027178 |
| *Pseudomonadaceae* | 166.4812 | 0.286513 | 0.147584 | 0.142084 |
| *Pseudonocardiaceae* | 103.1054 | -0.48027 | 0.186006 | 0.032744 |
| *Rhizobiaceae* | 540.1628 | 0.190274 | 0.061999 | 0.010739 |

| | | | | |
|---|---|---|---|---|
| *Rhodomicrobiaceae* | 222.2028 | 0.566638 | 0.102377 | 1.15E-06 |
| *Roseiflexaceae* | 246.7347 | 0.436395 | 0.133666 | 0.006443 |
| *Rubritaleaceae* | 404.6714 | 0.07716 | 0.106963 | 0.567085 |
| *Saccharimonadaceae* | 214.7912 | 0.164709 | 0.105573 | 0.247343 |
| *Solirubrobacteraceae* | 153.0475 | 0.174746 | 0.163668 | 0.432822 |
| *Sphingobacteriaceae* | 319.3659 | -0.48343 | 0.119487 | 0.000586 |
| *Spirosomaceae* | 163.2731 | -0.09857 | 0.12104 | 0.521632 |
| *Streptomycetaceae* | 189.924 | -0.12651 | 0.11795 | 0.432822 |
| *Xanthobacteraceae* | 360.5323 | -0.25971 | 0.072029 | 0.002396 |
| **Comparison – Rhizosphere-Endosphere** | | | | |
| **OTU** | **Base Mean** | **log2 Fold Change** | **lfcSE*** | **padj** |
| *Bacillaceae* | 669.8586 | -0.24338 | 0.093574 | 0.02817 |
| *Bacteriovoracaceae* | 211.6517 | -0.38527 | 0.16022 | 0.0426 |
| *Burkholderiaceae* | 1560.524 | -0.05437 | 0.071256 | 0.556862 |
| *Caulobacteraceae* | 161.1426 | -0.32272 | 0.179428 | 0.126463 |
| *Chitinophagaceae* | 2810.665 | 0.334696 | 0.09971 | 0.003625 |
| *Gaiellaceae* | 616.729 | 0.668651 | 0.14625 | 5.37E-05 |
| *Geminicoccaceae* | 253.4938 | 0.051528 | 0.157662 | 0.835729 |
| *Gemmataceae* | 609.9269 | 0.873427 | 0.10693 | 1.56E-14 |
| *Haliangiaceae* | 412.6111 | 0.106914 | 0.097998 | 0.393264 |
| *Hymenobacteraceae* | 421.6195 | 0.664185 | 0.119973 | 6.19E-07 |
| *Ilumatobacteraceae* | 1702.794 | 1.175225 | 0.140792 | 6.99E-15 |
| *Microbacteriaceae* | 174.5953 | -0.56967 | 0.190141 | 0.01013 |
| *Micromonosporaceae* | 251.2053 | 0.040195 | 0.145248 | 0.839919 |
| *Mycobacteriaceae* | 430.8715 | 0.710153 | 0.17698 | 0.000429 |
| *Nitrosomonadaceae* | 687.0426 | 0.216746 | 0.141276 | 0.19838 |
| *Nitrospiraceae* | 613.7027 | 0.610044 | 0.137464 | 8.26E-05 |
| *Opitutaceae* | 1042.123 | 0.607463 | 0.131676 | 4.95E-05 |
| *Paenibacillaceae* | 959.6071 | 0.525368 | 0.079788 | 1.52E-09 |
| *Pedosphaeraceae* | 677.0895 | 0.550655 | 0.095806 | 2.26E-07 |
| *Planococcaceae* | 369.4337 | -0.28308 | 0.191611 | 0.214734 |
| *Pseudomonadaceae* | 195.1522 | 0.386658 | 0.234509 | 0.165314 |
| *Pseudonocardiaceae* | 222.4399 | 0.516128 | 0.159337 | 0.005211 |
| *Rhizobiaceae* | 385.9288 | -0.45277 | 0.114062 | 0.00048 |
| *Rhodomicrobiaceae* | 100.4318 | -0.77128 | 0.17287 | 8.13E-05 |
| *Roseiflexaceae* | 144.6883 | -0.43948 | 0.214584 | 0.08816 |
| *Rubritaleaceae* | 366.9267 | -0.17432 | 0.175362 | 0.444729 |
| *Saccharimonadaceae* | 209.715 | 0.035154 | 0.126667 | 0.839919 |
| *Solirubrobacteraceae* | 140.2452 | -0.04875 | 0.182615 | 0.839919 |
| *Sphingobacteriaceae* | 599.1594 | 0.360183 | 0.106568 | 0.003625 |
| *Spirosomaceae* | 187.2959 | -0.00022 | 0.28912 | 0.999402 |
| *Streptomycetaceae* | 371.4646 | 0.613746 | 0.142608 | 0.00014 |

| | | | | |
|---|---|---|---|---|
| *Xanthobacteraceae* | 370.2052 | -0.33994 | 0.144082 | 0.04694 |
| *lfsSE= log2 Fold Change Standard Error | | | | |

Supplementary Table 14. DESeq2 outputs stem elongation/senescent plants- Significantly differentially abundant bacterial taxa

| **Comparison – Senescent endosphere/Stem elongation endosphere** | | | | |
|---|---|---|---|---|
| **OTU** | **Base Mean** | **log2 Fold Change** | **lfcSE*** | **padj** |
| *Bacillaceae* | 357.5098 | -0.81595 | 0.241208 | 0.004485 |
| *Bacteriovoracaceae* | 25.6759 | -1.66011 | 0.594842 | 0.019471 |
| *Burkholderiaceae* | 3318.41 | -1.25399 | 0.238716 | 1.66E-06 |
| *Caulobacteraceae* | 541.1063 | -0.79572 | 0.360076 | 0.072479 |
| *Chitinophagaceae* | 870.3272 | 0.176643 | 0.254357 | 0.587213 |
| *Devosiaceae* | 528.9238 | -1.11192 | 0.28085 | 0.000579 |
| *Gaiellaceae* | 68.69234 | 0.214398 | 0.45143 | 0.69004 |
| *Geminicoccaceae* | 38.29272 | 0.370222 | 0.423023 | 0.515508 |
| *Haliangiaceae* | 47.10503 | 0.248007 | 0.470804 | 0.657529 |
| *Hymenobacteraceae* | 19.25489 | -0.84456 | 0.475021 | 0.157109 |
| *Ilumatobacteraceae* | 160.0597 | 0.364665 | 0.301336 | 0.395293 |
| *Microbacteriaceae* | 656.3924 | -0.31577 | 0.245576 | 0.354469 |
| *Micromonosporaceae* | 1086.754 | -1.17346 | 0.357779 | 0.005467 |
| *Mycobacteriaceae* | 84.1459 | -0.47063 | 0.259804 | 0.149081 |
| *Nitrosomonadaceae* | 43.46863 | 0.355034 | 0.473786 | 0.567054 |
| *Nitrospiraceae* | 33.82162 | 0.659862 | 0.322814 | 0.090987 |
| *Opitutaceae* | 22.22379 | -0.56716 | 0.489488 | 0.407432 |
| *Paenibacillaceae* | 431.831 | -1.74822 | 0.248606 | 5.09E-11 |
| *Pedosphaeraceae* | 18.52983 | 0.736093 | 0.466216 | 0.219935 |
| *Planococcaceae* | 59.45307 | -0.31161 | 0.280003 | 0.407432 |
| *Promicromonosporaceae* | 1086.754 | -1.17346 | 0.357779 | 0.005467 |
| *Pseudomonadaceae* | 304.2958 | -0.94093 | 0.341331 | 0.020137 |
| *Pseudonocardiaceae* | 878.523 | -0.10126 | 0.262648 | 0.73668 |
| *Rhizobiaceae* | 1302.42 | 0.255903 | 0.231463 | 0.407432 |
| *Rhodomicrobiaceae* | 5.325793 | 0.713473 | 0.635217 | 0.407432 |
| *Roseiflexaceae* | 104.1034 | -1.21258 | 0.403322 | 0.010405 |
| *Rubritaleaceae* | 238.5271 | -0.68881 | 0.318014 | 0.075784 |
| *Saccharimonadaceae* | 613.9465 | 0.60443 | 0.56084 | 0.41666 |
| *Solirubrobacteraceae* | 40.43118 | -0.13417 | 0.411122 | 0.775156 |
| *Sphingobacteriaceae* | 1977.09 | -1.36091 | 0.330803 | 0.000324 |
| *Spirosomaceae* | 815.2618 | -1.9214 | 0.301822 | 3.23E-09 |
| *Streptomycetaceae* | 7007.769 | -2.15252 | 0.314852 | 1.62E-10 |
| *Xanthobacteraceae* | 424.1464 | 0.19374 | 0.240251 | 0.554194 |
| *lfsSE= log2 Fold Change Standard Error | | | | |

Supplementary Table 15. DESeq2 outputs Significantly differentially abundant fungal taxa

| **Comparison - Bulk Soil-Rhizosphere Senescent** | | | | |
|---|---|---|---|---|
| **OTU** | **Base Mean** | **log2 Fold Change** | **lfcSE*** | **padj** |
| *Ambisporaceae* | 136.0381 | -7.31509 | 1.504427 | 3.60E-05 |
| *Chaetosphaeriaceae* | 583.7226 | -4.90418 | 1.272226 | 0.000891 |
| *Cladosporiaceae* | 1751.722 | 0.352167 | 0.865945 | 0.861533 |
| *Erythrobasidiaceae* | 88.04139 | -0.04082 | 0.908122 | 0.964151 |
| *Hypocreales Incertae sedis* | 11149.53 | -0.45642 | 0.716685 | 0.774593 |
| *Mortierellaceae* | 3301.221 | -2.72388 | 0.842895 | 0.004771 |
| *Sporidiobolaceae* | 535.6329 | 3.439894 | 0.856549 | 0.000612 |
| *Sydowiellaceae* | 524.9519 | -0.50637 | 0.762741 | 0.774593 |
| *Parmeliaceae* | 40.84577 | 5.768819 | 1.221575 | 3.61E-05 |
| **Comparison – Rhizosphere-Endosphere Senescent** | | | | |
| **OTU** | **Base Mean** | **log2 Fold Change** | **lfcSE*** | **padj** |
| *Ambisporaceae* | 1.053667 | 1.051434 | 2.149638 | 0.756691 |
| *Chaetosphaeriaceae* | 43.17347 | 2.16618 | 0.73166 | 0.019034 |
| *Cladosporiaceae* | 438.7817 | -1.79186 | 0.722699 | 0.049133 |
| *Erythrobasidiaceae* | 19.80491 | -4.76126 | 1.234989 | 0.001089 |
| *Hypocreales Incertae sedis* | 2296.368 | -2.83945 | 0.668941 | 0.000339 |
| *Mortierellaceae* | 225.3781 | -1.99924 | 0.697942 | 0.021581 |
| *Parmeliaceae* | 15.73176 | 1.890016 | 1.11563 | 0.233124 |
| *Sporidiobolaceae* | 191.0621 | -3.80797 | 0.717967 | 3.52E-06 |
| *Sydowiellaceae* | 117.4911 | -1.88265 | 0.696616 | 0.030472 |
| **Comparison – Bulk Soil-Rhizosphere Stem Elongation** | | | | |
| *Ambisporaceae* | 1.126446 | 0.297184 | 1.528897 | 0.895664 |
| *Chaetosphaeriaceae* | 17.49084 | -0.70025 | 1.581208 | 0.895664 |
| *Cladosporiaceae* | 499.0058 | -0.33263 | 0.773925 | 0.895664 |
| *Erythrobasidiaceae* | 266.2979 | 2.846739 | 1.621902 | 0.369093 |
| *Hypocreales Incertae sedis* | 8407.459 | -1.26002 | 0.733516 | 0.369093 |
| *Mortierellaceae* | 3348.64 | 3.034097 | 0.857525 | 0.017323 |
| *Sporidiobolaceae* | 979.9934 | 0.496755 | 0.954706 | 0.895664 |
| *Sydowiellaceae* | 810.7476 | -1.01088 | 0.795848 | 0.461721 |
| *Parmeliaceae* | 946.0702 | 0.948964 | 0.83265 | 0.497266 |
| **Comparison – Rhizosphere-Endosphere Stem Elongation** | | | | |
| *Ambisporaceae* | 3.5 | 2.584955 | 1.466257 | 0.136337 |
| *Chaetosphaeriaceae* | 6.666667 | -2.23704 | 1.439501 | 0.168246 |
| *Cladosporiaceae* | 211.6667 | -3.02444 | 1.295549 | 0.053817 |
| *Erythrobasidiaceae* | 191.5 | -5.43269 | 1.6628 | 0.004345 |
| *Hypocreales Incertae sedis* | 2356.833 | -2.30279 | 0.561104 | 0.000314 |
| *Mortierellaceae* | 2594.5 | -6.9257 | 0.786027 | 3.47E-17 |
| *Parmeliaceae* | 2737.5 | 2.135562 | 0.52331 | 0.000314 |

| | | | | |
|---|---|---|---|---|
| *Sporidiobolaceae* | 525.3333 | -6.51894 | 1.131744 | 1.18E-07 |
| *Sydowiellaceae* | 239.1667 | -3.20945 | 0.839198 | 0.000734 |
| *lfsSE= log2 Fold Change Standard Error | | | | |

Supplementary Table 16. DESeq2 outputs Significantly differentially abundant fungal taxa

| **Comparison - Endosphere Senescent-Endosphere Stem Elongation** | | | | |
|---|---|---|---|---|
| **OTU** | **Base Mean** | **log2 Fold Change** | **lfcSE*** | **padj** |
| *Parmeliaceae* | 2232.5 | -9.80089 | 0.980524 | 4.94E-22 |
| *Leotiaceae* | 2225.5 | -12.1196 | 1.281843 | 5.02E-20 |
| *Myxotrichaceae* | 524.5 | -7.1502 | 1.058346 | 1.47E-10 |
| *Clavicipitaceae* | 4820.833333 | -3.95072 | 1.096925 | 0.0014004 |
| *Chaetosphaeriaceae* | 26.66666667 | 4.450025 | 1.070374 | 0.0001663 |
| **Comparison – Bulk Soil-Rhizosphere all samples (compost, agricultural pot, 50:50 mix and field stem elongation)** | | | | |
| **OTU** | **Base Mean** | **log2 Fold Change** | **lfcSE*** | **padj** |
| *Australiascaceae* | 736.0431973 | -0.22788 | 0.803202 | 0.8926117 |
| *Glomerellaceae* | 2480.256112 | -0.23093 | 0.96327 | 0.8926117 |
| *Hypocreales Incertae sedis* | 6355.970558 | -1.04999 | 0.731854 | 0.509278 |
| *Leotiaceae* | 1357.387391 | -1.72508 | 0.760808 | 0.2009295 |
| *Mortierellaceae* | 1382.193766 | 3.010589 | 0.87225 | 0.0119848 |
| **Comparison – Rhizosphere-Endosphere all samples (compost, agricultural pot, 50:50 mix and field stem elongation)** | | | | |
| *Australiascaceae* | 616.0318099 | -3.69862 | 0.722446 | 6.58E-06 |
| *Glomerellaceae* | 2071.880172 | -3.3994 | 0.928719 | 0.0009847 |
| *Hypocreales Incertae sedis* | 3357.081144 | -3.56054 | 0.733033 | 1.59E-05 |
| *Leotiaceae* | 1491.808491 | 3.60561 | 0.749014 | 1.59E-05 |
| *Mortierellaceae* | 1463.011582 | -3.95723 | 1.01644 | 0.0004576 |
| *lfsSE= log2 Fold Change Standard Error | | | | |
